# Current sequence-based models capture gene expression determinants in promoters but mostly ignore distal enhancers

**DOI:** 10.1101/2022.09.15.508087

**Authors:** Alexander Karollus, Thomas Mauermeier, Julien Gagneur

## Abstract

**Background:** The largest sequence-based models of transcription control to date have been obtained by predicting genome-wide gene regulatory assays across the human genome. This setting is fundamentally correlative, as those models are exposed during training solely to the sequence variation between human genes that arose through evolution, questioning the extent to which those models capture genuine causal signals.

**Results:** Here we confront predictions of state-of-the-art models of transcription regulation against data from two large-scale observational studies and five deep perturbation assays. The most advanced of these sequence-based models, Enformer, by and large captures causal determinants of human promoters. However, models fail to capture the causal effects of enhancers on expression, notably in medium to long distances and particularly for highly expressed promoters. More generally, the predicted impact of distal elements on gene expression predictions is small and the ability to correctly integrate long-range information is significantly more limited than the receptive fields of the models suggest. This is likely caused by the escalating class imbalance between actual and candidate regulatory elements as distance increases.

**Conclusions:** Our results suggest that sequence-based models have advanced to the point that in-silico study of promoter regions and promoter variants can provide meaningful insights and we provide practical guidance on how to use them. Moreover, we foresee that it will require significantly more and particularly new kinds of data to train models accurately accounting for distal elements.

## Introduction

Regulatory regions in the genome encode instructions determining gene product abundance in response to developmental and environmental cues encode instructions determining gene product abundance in response to developmental and environmental cues. Inherited or acquired genetic alterations in these regulatory regions can result in the dysregulation of gene expression, which ultimately can cause a variety of diseases. Accordingly, models which can reliably predict gene expression directly from sequence would not only be of scientific interest but could potentially find many uses in the design of personalized diagnoses and treatments.

Currently, no sequence-based model is capable of holistically accounting for all stages of gene expression from transcription initiation to protein degradation and can thus predict the abundance of each processed protein isoform in any given cellular context. However, deep learning models have recently been proposed which - at least in theory - can predict measures of RNA abundance directly from arbitrary input sequences centered on a gene of interest for large sets of human cell types and tissues. The focus of this study is to understand how successful these models are. For the sake of simplicity, we will follow the convention in the literature and use RNA abundance and gene expression synonymously, even if the former is only an imperfect proxy of the latter [1,2].

We will study Xpresso [3], a cell-type agnostic model which predicts gene expression from a small sequence window around the transcription start site (TSS), Basenji1 [4] and 2 [5], deep convolutional models which were trained on many genome-wide assays and cell lines of the ENCODE project [6,7] and use about 40 kilobases (kb) of context, and Expecto [8], a linear model trained on top of the deep convolutional model DeepSea [9], itself trained to predict ENCODE genome-wide assays. Moreover, we will investigate the performance of the largest model trained to date, Enformer [10], a deep transformer [11] with ∼250 million trainable parameters - an order of magnitude more than preceding models. As input, Enformer gets a 196 kb long sequence and predicts the value of 5,313 different ENCODE tracks in bins of 128bp. These tracks include chromatin-immunoprecipitation signal for hundreds of transcription factors, DNase footprinting (DNase), which measures genome accessibility, and cap-analysis of gene expression (CAGE) measurements for hundreds of cellular contexts (defined as combinations of cell lines and treatments). For this study, the CAGE predictions are the most relevant, as these provide the sought-after measure of RNA abundance.

As Avsec et al. [10] convincingly show in their paper, Enformer provides unparalleled performance when predicting CAGE signal of held-out test genes. At least in some cell types, the model is close to experimental accuracy. However, such aggregate measures of performance carry significant caveats. The main issue is that all the models we named were trained using the sole genetic diversity available across the human and mouse genomes. This setting is fundamentally correlative, as genomic sequences of any organism are not a random sample from the space of possible sequences but rather have been selected and co-evolved over millions of years of evolution. Thus, it is unclear to what extent the models have learned and employ causal principles, rather than mere correlations, to make their predictions. If so, predictions may become very misleading when applied in a medical diagnostic context to interpret rare germline or somatic variants, or if the model is used to generate new mechanistic hypotheses.

Additionally, improved performance of a model on aggregate measures such as total explained variance does not provide insights into the reasons for the improvement. Enformer has more parameters than previous models, but also a much wider receptive field - i.e. the size of the sequence window it can integrate over to predict expression at a particular location. Is the wider receptive field the deciding factor for its improved performance? Understanding the actual source of performance improvements will help to design better models.

To address these questions, we conducted in-silico reproductions of two large-scale observational assays which measured gene expression in different tissues and stages of development and five deep perturbation assays, including designed massively parallel reporter assays for promoters and enhancers, CRISPRi enhancer-knockdown and saturation mutagenesis experiments. Moreover, we conducted two in-silico perturbation studies. In this way, we can probe in a targeted fashion to what extent a model actually makes use of particular regulatory elements in its expression predictions and whether it does so in a way consistent with experiments.

## Results

Evaluating Deep Models through in-silico reproduction of experiments

The overview figure (Fig 1) summarizes the different datasets we used in our study, which regulatory element(s) each dataset focuses on, and how we reproduced the experiment in-silico. Generally, replicating an experiment with a sequence-based model is not straightforward and requires three preparatory steps:

**Figure 1:**
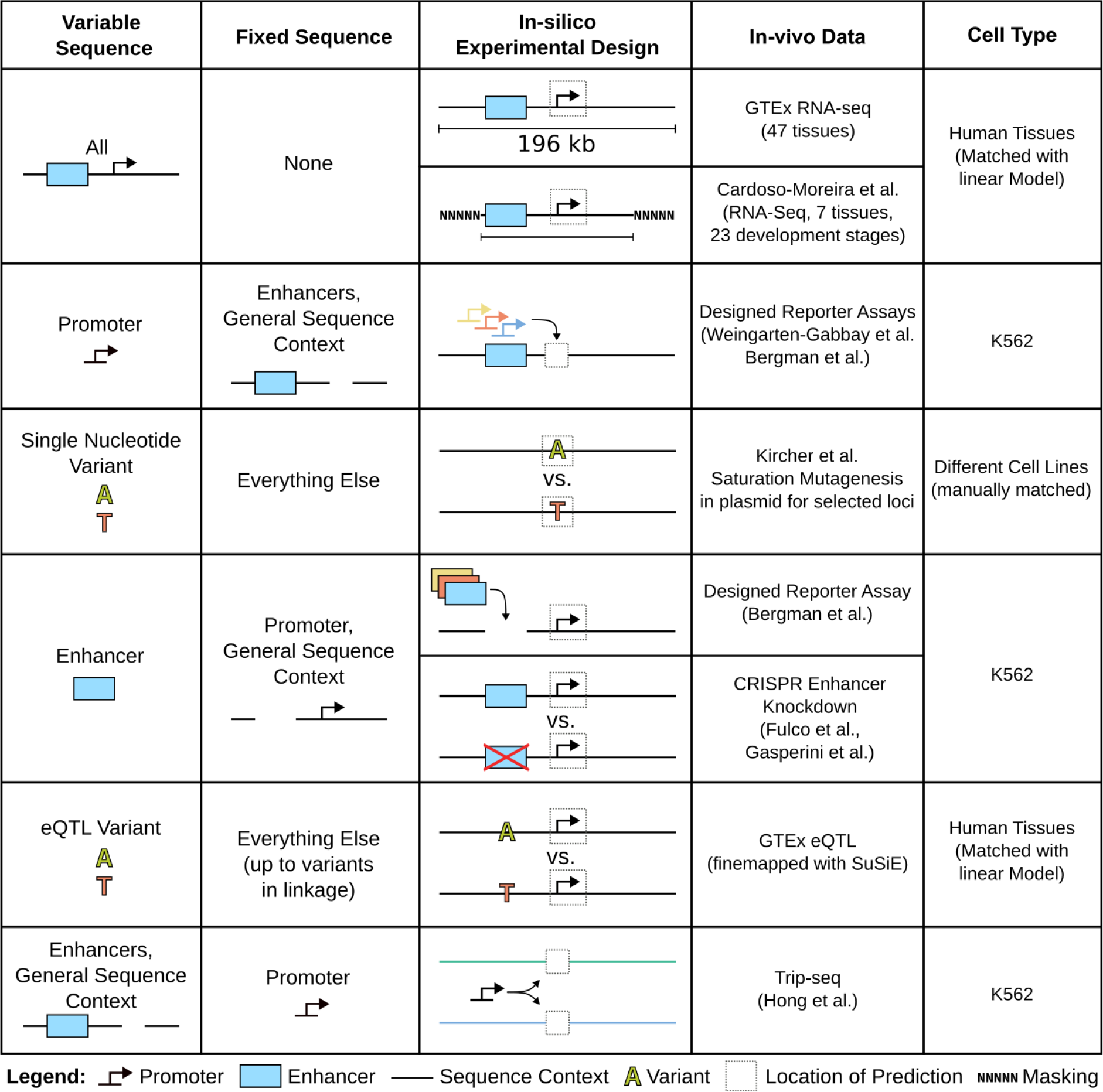
Overview of our in-silico experiments. To assess the generalization power of the models we performed analyses (rows), in which certain sequences varied (first column) while others were kept fixed (second column).

1. As many experiments involve some modification of the endogenous genome, we must construct in silico the correct sequences.
2. Most datasets do not report CAGE tracks but alternative measures of gene expression including gene-level RNA-sequencing read counts and reporter fluorescence. Hence, with the exception of Xpresso, which gives gene-level predictions, we need to decide for which transcription start site predictions should be made, i.e. from which bins to record the CAGE predictions.
3. Many experiments are conducted in cell types and tissues that do not exactly match those of ENCODE. Some matching or mapping between those cellular contexts must therefore be done.

The first step is the most intricate one. Several experiments we analyzed integrated sequences into the endogenous genome. Usually, these sequences consist of the regulatory element of interest, a reporter, post-transcriptional elements (e.g. chimeric introns and viral polyadenylation sites), and technical elements which facilitate the genomic integration (e.g. retrotransposon long terminal repeats) and sequencing. Many of these technical elements are highly artificial and thus unlike anything the deep models we consider will have seen in their training data. For this reason, we performed each replication twice, once with a faithful reproduction of the full insert and once with a minimal insert, consisting only of the regulatory element of interest and the reporter. Interestingly, almost always this minimal insert led to predictions that better correlated with the experiment. Therefore, We decided to report only the results from these minimal inserts. When possible, we avoided plasmid-based assays, as there is no way to represent a circular chromosome in the models we consider. For lack of alternatives, we made two exceptions, namely the Bergman et al. [12] promoter-enhancer compatibility study and the Kircher et al. [13] saturation mutagenesis study.

The second step is somewhat easier. For RNA-seq datasets (GTEx [14], Cardoso-Moreira et al. [15], GTEx eQTL [16]), we considered CAGE prediction for the TSS of the Ensembl [17] canonical transcript. This might not be the correct transcript in some tissues, but considerably reduces the number of predictions to be made. The only exception to this rule is the CRISPRi enhancer-knockout study, where we used the same TSS sites as Avsec et al. [10] If we did not know the location of the TSS, but we knew where the core promoter was located (e.g. because the experiment involves integrating promoters at arbitrary locations, then we took the prediction around the promoter midpoint. For the Kircher et al. saturation mutagenesis study, we took the prediction at the variant as done previously ([10]).

For the third step, all but three of the experiments we considered were done in K562 cells. In these cases, we simply took the K562 CAGE track. For the studies done using RNA-seq of human tissue biosamples, we fitted for each tissue a simple linear model (ridge regression) predicting RNA abundance from all ENCODE CAGE tracks. We held out the Enformer/Basenji2 test-set genes while fitting these regressions and evaluated only on this held-out data. These fitted regressions were also used to analyze GTEx eQTL. For the Kircher et al. saturation mutagenesis data, we used a manual matching procedure.

Once these preparatory steps are completed, we can run the sequence-based model on the constructed sequences and collect the relevant predictions. Here it is important to note that many of these models are sensitive to small changes in the input (e.g. small shifts), particularly if regulatory elements fall directly on bin boundaries. To mitigate this, we computed each prediction six times - for both strands and with small offsets respectively - and took the average. We also always summed predictions over three neighboring bins. The same technique was used by Avsec et al. [10], presumably for the same reason.

### Deep Models, particularly Enformer, provide very accurate predictions of gene expression in human tissues and during human development

Avsec et al. [10] already provide ample evidence that Enformer can predict the expression of endogenous genes very well. However, these validations were mostly done on ENCODE, which is highly enriched for cancer cell lines that may provide an imperfect proxy of in-vivo expression. Thus, to provide a slightly more complete picture, we benchmarked the model on two additional RNA-Seq datasets: GTEx [14], which measured gene expression in 49 human tissues, and Cardoso-Moreira et al. [15] which measured gene expression in 7 tissues for 23 stages of development - from 4 weeks post-conception to senescence.

In both datasets, Enformer performs very well at predicting the between-gene variation (Fig 2A). For the median GTEx tissue, predictions and measurements correlate with r = 0.79. Moreover, Enformer consistently outperforms both Xpresso and Basenji2. Adding the tissuespecific exon-intron ratio of each gene - a (weak) proxy of RNA stability - as an additional predictor improves the performance on GTEx. This suggests that Enformer predictions could be augmented using a model that better captures post-transcriptional regulation. The median correlation in the Cardoso-Moreira et al. dataset is very similar (r = 0.77). Notably, performance is uneven across tissues and also degrades in later stages of development (Fig 2B, Fig S1). This might be due to varying data quality, the influence of environmental factors, because some tissues and stages of development intrinsically feature more complex regulation (e.g. testis [18]) or because they are not properly covered by the ENCODE cell lines Enformer was trained on.

**Figure 2:**
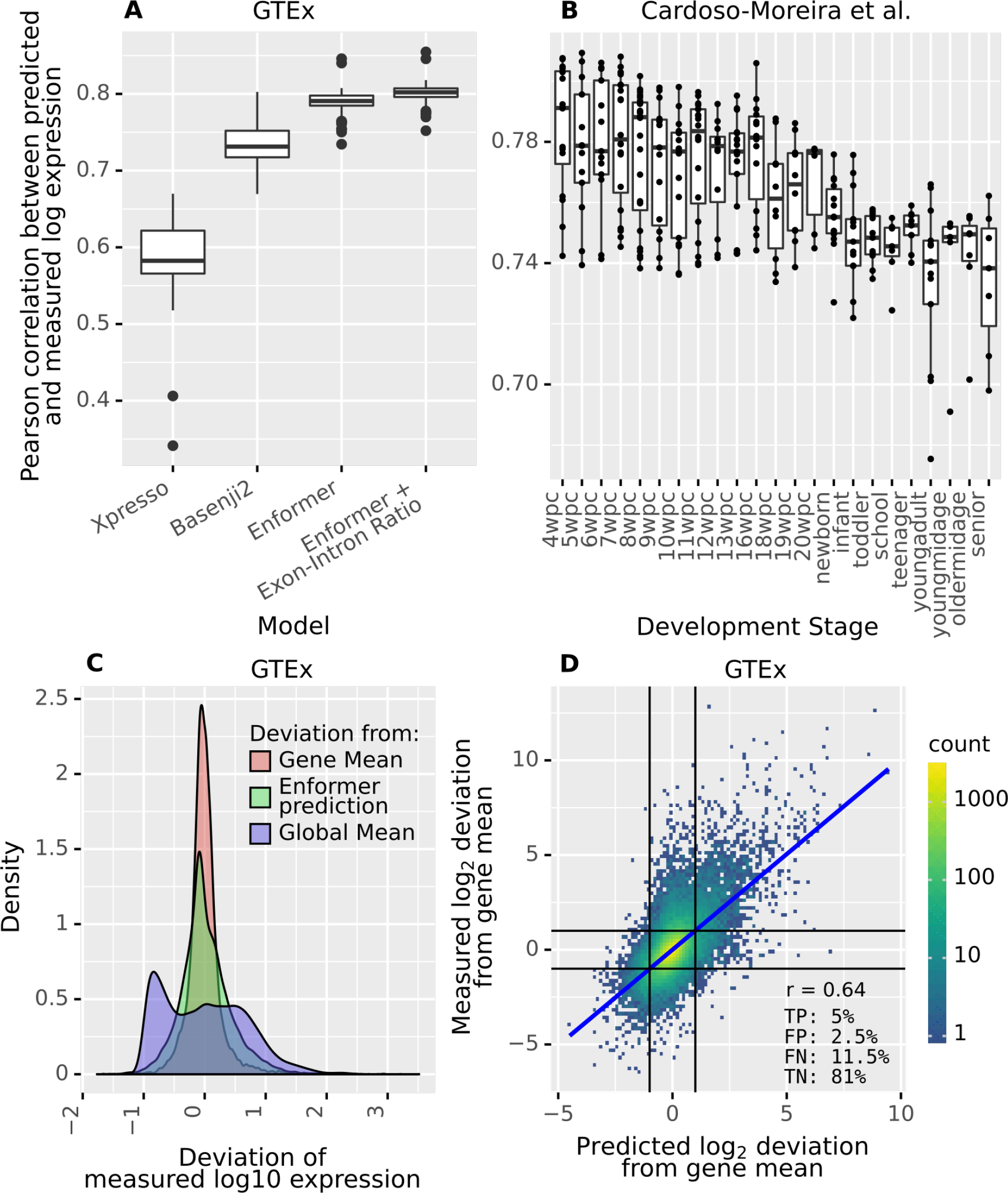
Enformer provides effective gene expression prediction for endogenous genes. **A)** Pearson correlation between predicted and measured log-transformed expression on GTEx tissues for different models. Enformer can predict endogenous RNA abundance, as measured in adult tissues (GTEx [14]), better than previous models. Adding the exon-intron ratio, a (weak) proxy of RNA half-life, as an additional predictor slightly improves performance. **B)** Same as A) for Enformer predictions on developmental samples (Cardoso-Moreira et al. [15] dataset). Enformer predicts endogenous gene expression very well overall yet somewhat worse for later stages of development. **C)** Distribution of deviations of GTEx measured log expression values from (1) the global mean (across genes and tissues, blue), (2) the gene mean (across tissues, red), and (3) the Enformer prediction (green). The first indicates overall variation in expression, the second indicates between-tissue variation and the third indicates the magnitude of errors of Enformer. Enformer accuracy is sufficient to explain much of the between-gene variation but not for the variation of genes between tissues. **D)** Measured between-tissue deviations of gene expression against prediction. Enformer predicts large between-tissue changes in expression reasonably well on average, but there is significant room for improvement. The numbers indicate the percentages of true positives (TP), false positives (FP), false negatives (FN), and true negatives (TN) when predicting 2-fold changes (black lines).

The above-mentioned correlations are computed across all genes, which span a large dynamic range. Specifically, the mean absolute deviation from the grand mean of log RNA abundance (across expressed genes and tissues) in the GTEx data is ∼4-fold (Fig 2C). In contrast, the mean absolute deviation to the mean per gene across tissue is only ∼1.5-fold. As Enformer has a mean-absolute error of about 2-fold (Fig 2C), it naturally struggles to predict smaller differences between tissues. Nevertheless, the correlation between measured and predicted log fold changes of genes between GTEx tissues is still remarkable (r = 0.64, Fig 2D, Basenji2: r = 0.54). This performance translates to a decent precision of 66% for a recall of 30% at predicting 2-fold changes between GTEx tissues.

Overall, we find that Enformer can predict endogenous RNA abundance very well and consistently outperforms previous models. This being said, when we compare the expression of different genes, we are comparing highly dissimilar regulatory sequences. These genes will generally have different promoters, different GC-content, different enhancers, and will be located in different chromosomal contexts. Thus, these aggregated results do not tell us which features of the sequence the model uses to make its predictions.

### Most of the receptive field has a very minor impact on Enformer gene expression predictions

Due to its wide receptive field (196kb), Enformer can theoretically account for the impact of regulatory elements up to a distance of 98 kb on either side of a TSS. Given our observations in the previous section, we wondered to what extent Enformer relies on these distal elements to correctly predict gene expression.

To this end, we created sequence windows of varying sizes centered on the TSS of genes by masking distal parts of the endogenous sequence (with “N” nucleotides). We then let Enformer predict the CAGE gene expression at these TSS for each window size. In this way, we can evaluate how the predictive power of Enformer on the GTEx [14] and the Cardoso-Moreira et al. [15] data changes when it can no longer use distal elements to inform its predictions.

Reassuringly, expanding the sequence window consistently improves the gene expression predictions (Fig 3A). However, Enformer already achieves substantial explained variance (about two-thirds of what is achieved with the full sequence) with only a tiny sequence window of 1001bp (∼0.5% of the total receptive field) around the TSS. Moreover, we face strong diminishing returns when adding additional sequence context. While expanding from 1kb to ∼40kb yields substantial improvements, the entire distal two-thirds of the sequence only added ∼1% of the total explained variation. Importantly, the improvement of Enformer against its predecessor Basenji2 appears consistent across all distances (Fig S2), with substantially better predictions even when Enformer can only access the same amount of sequence than Basenji2 (∼40kb), particularly for between-condition predictions.

**Figure 3:**
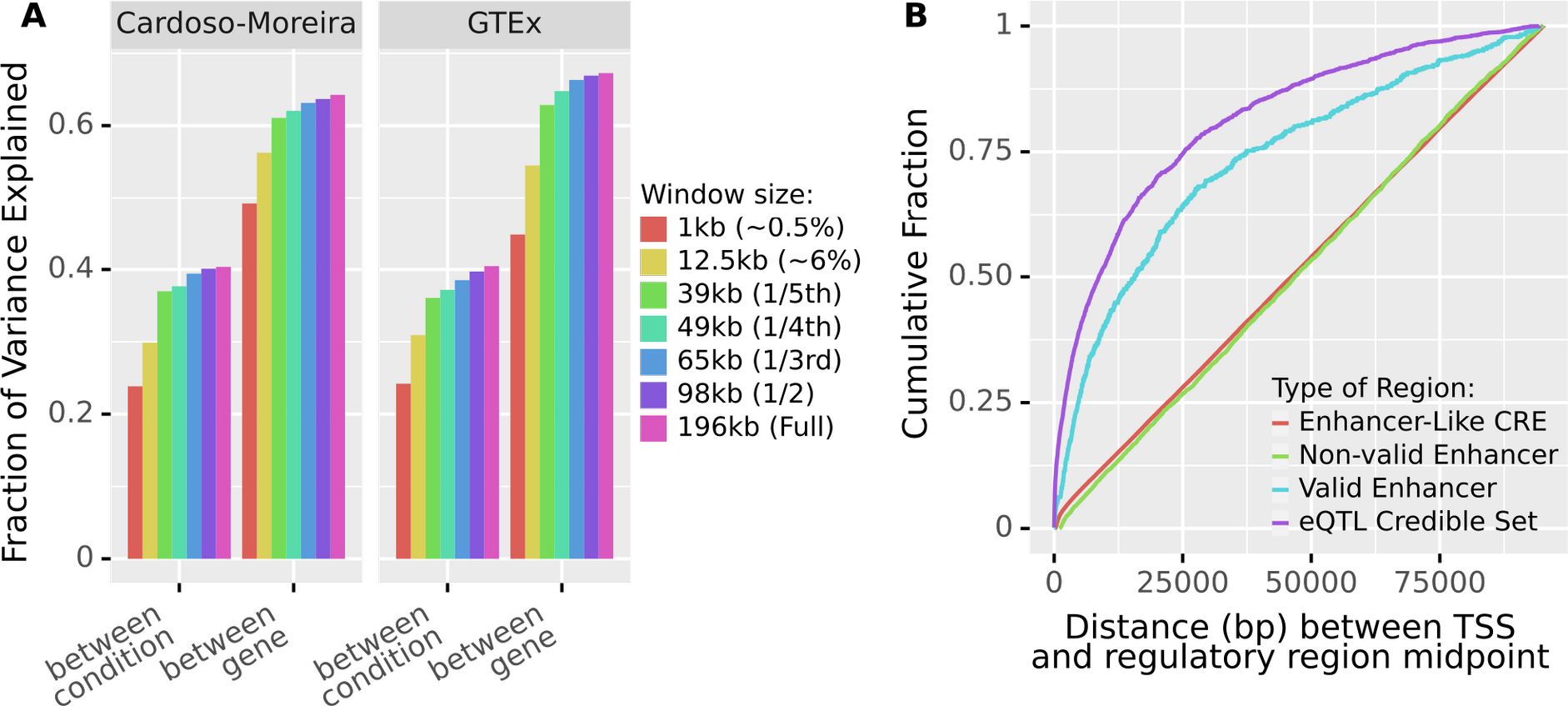
Enformer has very similar predictive power even if we severely restrict its input window, partially because most strong regulators are proximal. **A)** Fraction of variance in log-transformed expression, both between conditions and between genes, which Enformer can explain given varying amounts of sequence context. Values computed on Enformer held-out data. Most of the signal comes from the sequence immediately around the TSS, with the distal two-thirds of the input window contributing very little. **B)** Distribution of the distance within 98 kb of TSS of bona fide regulatory elements (eQTL, purple, and CRISPRi validated enhancers, blue) and candidate elements (ENCODE CRE with enhancer-like signal, red) and CRISPRi tested but not validated enhancers (green). Most bona fide regulatory elements lie close to their target gene whereas candidate elements are uniformly distributed. We only consider elements within 98kb of a TSS, i.e. within the Enformer receptive field.

In conclusion, Enformer extracts most of the signal to predict gene expression from promoter and promoter-proximal sequences, with sequences further than 30kb from the TSS having a negligible impact on its overall explained variance. Surprisingly, distal sequences also have little impact on predicting between-condition variation, where we would have expected relatively more influence from distal elements. What is not clear from this analysis is whether the seeming irrelevance of distal elements reflects biological reality.

### Most known strong regulators are located close to their target genes, leading to an extreme class imbalance at higher distances

Our previous results indicate that Enformer considers most distal sequences to have a negligible impact on gene expression at the TSS. We sought to examine the biological plausibility of this.

For this purpose, we computed the distance distribution of *bona fide* regulatory elements, specifically, eQTL [16] (which pass a series of filters, Methods) and CRISPRi validated enhancers [19,20]. Most of these *bona fide* regulatory elements are located close to the TSS of their target gene, in striking contrast to CRISPRi-tested but not validated candidate enhancers (Fig 3B). Because of the limited power of the underlying assays, we cannot necessarily conclude from this that most regulators of a gene will be proximal, but we can conclude that the majority of strong regulators (i.e. those with large, individual effects) will be proximal. These observations, consistent with previous studies [19,21,22], suggest that there is a biological basis for Enformer to attribute more importance to local sequences.

However, the distribution of bona fide regulatory elements seems to not be as imbalanced as the one implied by Enformer explained variance. Specifically, we see that the distal two-thirds of the sequence still contains ∼20% of known eQTL and ∼25% of validated enhancers. Interestingly, candidate regulatory elements from ENCODE (ENCODE SCREEN cis-Regulatory Elements with enhancer-like signature [23]) show a uniform distribution within 100 kb of TSS (Fig 3B). Thus, for a typical gene, we can usually find similar amounts of „ enhancer-like” sequences at every distance to the TSS. The result is that, as we move further from the TSS, the ratio of relevant regulatory elements to candidates (i.e. sequences which look reasonably similar to regulatory elements) will necessarily become very unfavorable very quickly. Thus, at higher distances, every long-range model will face an extreme class imbalance. Perhaps this class imbalance and the difficulty to distinguish enhancers targeting a given TSS from other enhancer-like elements is the cause of the apparent under-usage of distal regulatory sequences.

We next examined perturbation experiments in detail to determine whether individual regulatory elements contribute to Enformer predictions in a causal manner.

### Deep Models, particularly Enformer, correctly predict promoter strength and the impact of many promoter modifications

Most of the signal used by Enformer to predict gene expression seems to derive from the promoter and promoter-proximal elements. Thus we first sought to assess to what extent Enformer -and previous models -can predict the causal determinants of human promoters.

For this purpose, we performed an in-silico replication of the MPRA study by Weingarten-Gabbay et al. [24] who measured the impact of different endogenous and synthetic promoters on the expression of a reporter in K562 cells. In this study, 2274 164bp long sequence fragments corresponding to known human promoters and pre-initiation complex binding sites were inserted together with a fluorescence reporter at the AAVS1 viral integration site.

Generally, Enformer predicts the relative effects of those promoters very well (the Pearson correlation between measured and predicted log expression values is 0.68, Fig 4A). Moreover, Enformer outperforms all other models on this task.

**Figure 4:**
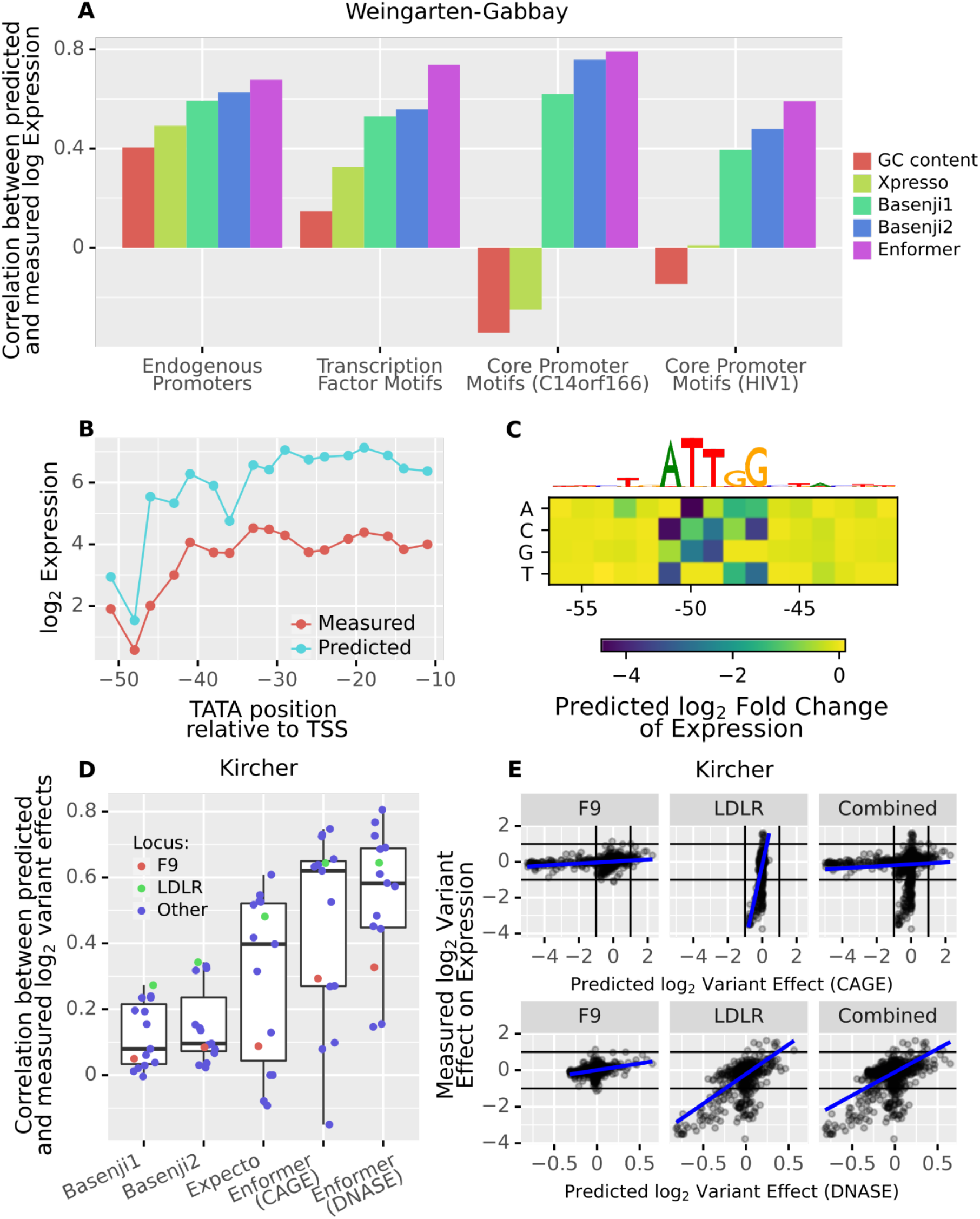
Enformer accurately predicts genetic perturbations of promoters. **A)** Correlation between model predictions and measurements across synthetic promoters of the Weingarten-Gabbay et al. [24] parallel reporter assay. Enformer outperforms preceding models, notably in targeted perturbation experiments (Transcription factor motif ablation and core promoter motif perturbations). **B)** Enformer can detect and often correctly interpret the impact on expression of subtle locational shifts of the TATA box in the RPLP0 promoter background. **C**) In-silico mutagenesis analysis suggests that the large drop of expression around position -50 likely from panel B is due to the disruption of a CAT-box at this location, rather than the positional preference of the TATA box. **D)** Pearson correlation between predicted and observed variant effects for the different loci tested by Kircher et al. [13] Enformer outperforms other models for most loci. **E)** Measured vs. predicted variant effects for two loci (F9, LDLR) individually and then both loci combined for CAGE (top) and DNase (bottom). The CAGE predictions appear to be miscalibrated across those two loci.

Weingarten-Gabbay et al. additionally constructed synthetic sequences to measure how individual promoter elements affect its overall strength. In one experiment, they inserted 133 different TF binding sites in two different backgrounds. We find that Enformer predicts the expression impact of these different TF motifs very well in both backgrounds (r = 0.75 and 0.73 respectively) and outperforms all other models (Fig 4A). We conclude that Enformer not only recognizes these motifs but also generally correctly determines whether they act as repressors, activators, or neither in K562 cells.

In yet another experiment, Weingarten-Gabbay et al. tested many combinations of six known core promoter motifs (including the TATA box and the initiator) in five backgrounds. Enformer predicts the impact of these perturbations in two backgrounds (r = 0.78 and 0.59 respectively, Fig S3), but no significant correlation between predictions and measured values was found for the other three backgrounds (but this is also true for the other models tested). On these two backgrounds, Enformer once again outperforms the competition (Fig 4A).

Additionally, Weingarten-Gabbay et al. measured the effect of shifting the position of a TATA box in four different backgrounds. In two out of four backgrounds, Enformer predictions show significant correlations with the measured effects (Fig S4). In the RPLP0 background, in particular, the correlation is almost perfect (r = 0.9, Fig 4B). Given that Enformer pools the sequence into 128bp bins, it is quite impressive that it is sensitive to such small shifts. Note, however, that Basenji1 and Basenji2 -but not Xpresso - deliver similar predictions on this task.

We wondered whether the large drop in both measured and predicted expression when the TATA was placed around position -50 was a result of the position preference of the TATA or due to another factor. As Weingarten-Gabbay et al. do not discuss this, we sought to use Enformer to answer this question. For this purpose, we first shifted a neutral k-mer (NNNNNNNN) through the sequence instead of the TATA. We find that this had little impact, except again around position -50 (Fig S5). We then performed an in-silico mutagenesis around this position and found that the expression prediction was most sensitive to the bases ATTGG (Fig 4C). This Enformer-based analysis provides a biologically plausible hypothesis, namely that the drop in expression is caused by a disruption of a CAT-box rather than due to a positional preference of the TATA box.

Lastly, we reanalyzed the Kircher et al. [13] saturation mutagenesis data which was already used as validation in the Avsec et al. [10] paper and additionally provided a comparison to Basenji1, Basenji2, and Expecto. In this experiment, short stretches of sequence from 15 loci (mostly promoters) were selected to serve as regulatory elements for a reporter gene in a plasmid. The authors then introduced almost every possible single nucleotide variant in these sequences and recorded their respective impacts on the expression of the reporter.

Consistent with Avsec et al., we found that observed and predicted relative variant effects (computed as log_2_ fold change of predicted expression at the variant, Methods) correlated well for most loci. Moreover, Enformer outperformed all other models (Fig 4D).

However, Enformer predictions appeared to be miscalibrated between loci (Fig S6). While a linear correlation is often present, the value of the slope varied drastically between loci. The problem is striking when comparing the F9 and LDLR loci, which were both measured and predicted in HepG2 (Fig 4E). As a consequence, while the individual correlations are significant for both loci (r = 0.3 and r = 0.64 respectively), the correlation drops substantially (r = 0.1) when pooling the data of the two loci. As those correlations are computed on log-transformed abundance, these observations imply that Enformer variant effect predictions only accurately reflect the experimental values up to a locus-specific exponential factor. This being said, it is unclear whether this discrepancy is due to Enformer or an artifact of the plasmid-based assay.

Interestingly, if we use accessibility (DNase) predictions as a proxy for RNA abundance predictions, then the effects are on average too small (a predicted log_2_ fold change of 0.65 corresponds roughly to a measured effect of 1), but this scaling factor is by and large consistent between loci (Fig 4E, Fig S7). Thus, for use cases such as genome-wide variant prioritization, DNase prediction score will likely be more useful.

Overall, we see that current sequence-based models, particularly Enformer, can match experimental data measuring the expression of different promoters in a controlled synthetic context very well. Moreover, it can predict the impact of different TF-binding sites, single nucleotide variants, and core promoter motifs, at least in some backgrounds. These analyses thus show that Enformer significantly captures genuine causal regulatory elements of the promoter.

### Enformer attributes considerably less importance to enhancers than experiments suggest

Having established that Enformer captures causal elements located in promoters, we next study to what extent it can correctly predict the effect of enhancers. Bergman et al. [12] assayed all combinations of one thousand 264 bp human promoter and one thousand 264 bp human enhancer fragments using a plasmid-based MPRA. A major result of this study was that transcriptional output appeared to be well modeled as the product of promoter strength and enhancer strength.

In agreement with our above-mentioned analysis of promoters, Enformer predicted the promoter strength of the Bergman et al. assay remarkably well (r = 0.81, Fig. 5A). In contrast, the predicted enhancer effect correlates poorly with the reported enhancer strength (r = 0.137, Fig. 5B). Moreover, promoters alone explained most of the Enformer predictions (90% predicted variation in log expression) but only half of the experimentally observed variation (54%, Fig. 5C).

**Figure 5:**
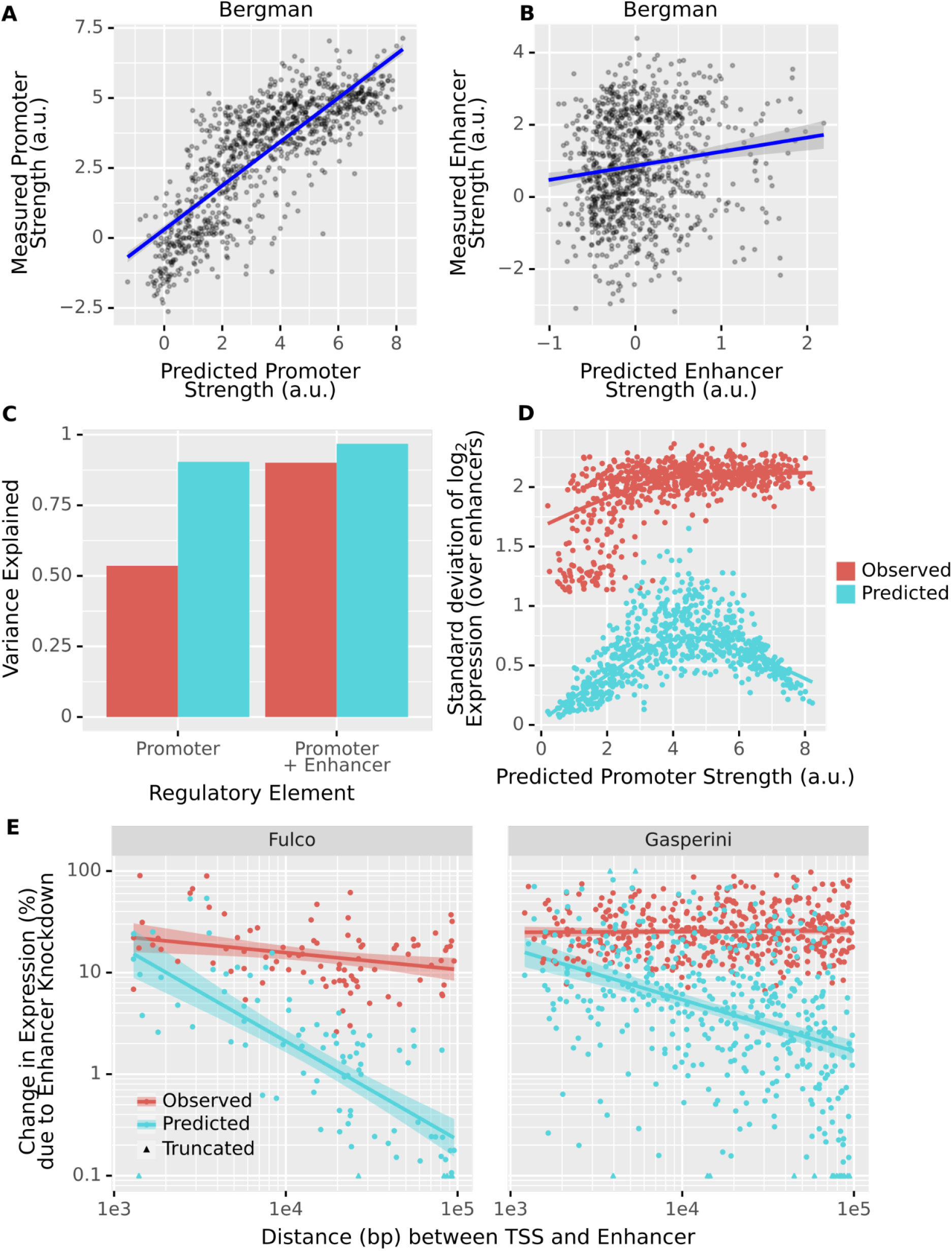
The predicted impact of enhancers, particularly distal enhancers, is significantly smaller than experiments suggest. **A)** Predicted promoter strength vs measured promoter strength. These strengths are determined by fitting linear models to the data/predictions **B)** Predicted enhancer strength vs measured enhancer strength. **C)** The promoter can explain 54% of the variation in the measurements of Bergmann et al. [12], with enhancers explaining another 36%. However, 90% of the variation in Enformer predictions for the same sequences is driven by the promoter alone. Thus, at least in a plasmid context, Enformer strongly underestimates the importance of the enhancer for determining gene expression. **D)** In Enformer, the predicted variation of expression induced by the enhancer also heavily depends on the promoter. Only promoters of intermediate predicted strength are sensitive to the choice of Enhancer. In the experimental data, strong and intermediate promoters show similar sensitivity. **C)** The measured and predicted changes in gene expression (expressed as an unsigned percentage) due to enhancer knockout as a function of the distance between the gene and the enhancer. Values < 0.1% and > 100% are truncated. Shown are only validated enhancer-gene pairs from Fulco et al. [20] and Gasperini et al. [19]. Enformer attributes significantly less effect to most validated enhancers than the experiments suggest. The effect is particularly strong for distal enhancers.

This is not to say that the enhancers never matter in the predictions. If we plot, for each promoter, the standard deviation in predicted log expression induced by the different enhancers, we notice that the choice of enhancer does sometimes matter, but only for promoters of intermediate predicted strength (Fig 5D). Promoters that Enformer considers very strong seem to basically override the enhancer. In the experimental data, we do not observe such a pattern.

Overall, this analysis indicates that Enformer inadequately accounts for the impact of enhancers on gene expression and cannot predict their measured average effects. However, one limitation of the Bergman et al. study is that it is based on a plasmid construct, which Enformer was not trained to handle and may not reflect endogenous gene regulation. Moreover, in this assay, enhancers are placed very close to the promoter and are part of the transcript, which may interfere with co-and post-transcriptional mechanisms.

Given these concerns, we wondered whether the results described above generalize when we apply Enformer to enhancer-promoter pairs in the endogenous genome. To find such pairs, we use the enhancer knockdown screens performed by Gasperini et al. [19] and Fulco et al. [20] In these screens, CRISPR interference (CRISPRi) is used to perturb enhancers and then the corresponding impact on gene expression is measured.

To replicate the CRISPRi knockdown experiments in the model, we used an in-silico mutagenesis (ISM) approach. Specifically, we computed the change in predicted expression at the gene TSS upon shuffling the enhancer sequence (Methods), assuming that a shuffled enhancer sequence should generally be non-functional. To account for random noise introduced by shuffling, we repeated the procedure 25 times and recorded the average impact.

We find that the model sometimes attributes plausibly large effects to enhancers, particularly if they are promoter-proximal. However, as the distance between the TSS and the enhancer increases, the model very quickly begins to excessively discount the importance of the enhancer for gene expression (Fig 5E). This discounting scales proportionally to the inverse of distance, which is not what is observed for validated enhancers. This is a strong decay, which is already substantial at a distance of only 10 kb (2% for Fulco et al, 5% for Gasperini et al., Fig 5E).

In the respective assays, knocking down the median validated enhancer (n = 522) has a measured impact on expression of ∼20%. According to Enformer, this median effect is only ∼4%. Moreover, for ∼60% of genes with a validated enhancer (n = 385), none of the tested candidates (whether validated or not, n = 2070) has an impact on expression that exceeds 10% (Fig S8).

As for the Bergman et al. [12] data, Enformer predicts smaller effects of enhancer knockdown for genes with high predicted basal expression (i.e. the predicted expression after knockdown, Fig S9). Whether or not a similar relationship exists in the experimental data is difficult to say, because the underlying assays have higher power to detect smaller effects for more highly expressed genes.

Our results do not contradict those of Avsec et al. [10], who showed with the same data that Enformer performs reasonably well at prioritizing validated enhancer-promoter pairs. In fact, on average, Enformer does attribute somewhat larger effects to validated than to non-validated enhancers, after controlling for distance and basal expression of the promoter (Fig S10). However, even when it correctly links enhancer and gene, this impact is usually small. If we rank enhancers by their effect, then the threshold for ∼50% recall of validated enhancers is a ∼3% predicted effect (Fig S11). Moreover, we here defined the enhancer impact in relative terms, i.e. as percentage change, as we consider this to be the biologically plausible metric. However, it is possible to achieve a slightly better classification in enhancer prioritization using the predicted absolute change as in the original study (Fig S12). This is because, in contrast to the predicted percentage change, which as we noted above, decreases with promoter strength, the absolute change increases proportionally to the predicted basal expression (Fig S13). Since predicted expression correlates well with actual expression, this metric thus privileges highly expressed genes. This likely adds to the classification performance because highly expressed genes mechanically will tend to have more validated regulatory elements, due to the limited power of the underlying assay (Fig S14). We, therefore, believe that the predicted absolute change delivers somewhat inflated performance.

Overall, the model strongly underestimates the effect of known enhancers, particularly if they are distal to the TSS, for gene expression regulation.

### Most distal eQTLs do not have a meaningful impact on expression predictions

Given our results for enhancers, we next asked whether similar observations also apply for expression Quantitative Trait Loci (eQTLs).

To test this, we let Enformer predict the impact on gene expression of GTEx eQTLs [16]. We applied a set of filters to exclude eQTLs that may act post-transcriptionally, have unclear associated TSS, and for which Enformer cannot be straightforwardly applied (Methods). A fine-mapping method was applied to associate each eQTL with a credible set of likely-causal variants and account for linkage (SuSie [25,26]). We then computed the predicted impact on expression at the canonical TSS of the target gene for all variants in the credible set. We defined the predicted eQTL effect as the maximum of these individual impacts.

As for the enhancer analysis, Enformer predicted effects of eQTLs were large when close to the TSS but decayed quickly and proportionally to the inverse of distance (Fig 6A). This decay was extreme, showing an average predicted effect of less than 1% from 1kb on. We do not see such decay in the measured normalized effect sizes of these eQTLs (Fig 6B). Although the GTEx normalized effect sizes do not correspond to expression fold-changes due to non-linear and gene-specific data preprocessing [16], a decreasing relationship with distance would be expected if Enformer’s predictions were correct. Moreover, Enformer predicts for those eQTLs surprisingly tiny effects (median = 0.5% change). This gives further evidence that the model strongly underweights the causal effect of distal regulatory elements on gene expression.

**Figure 6:**
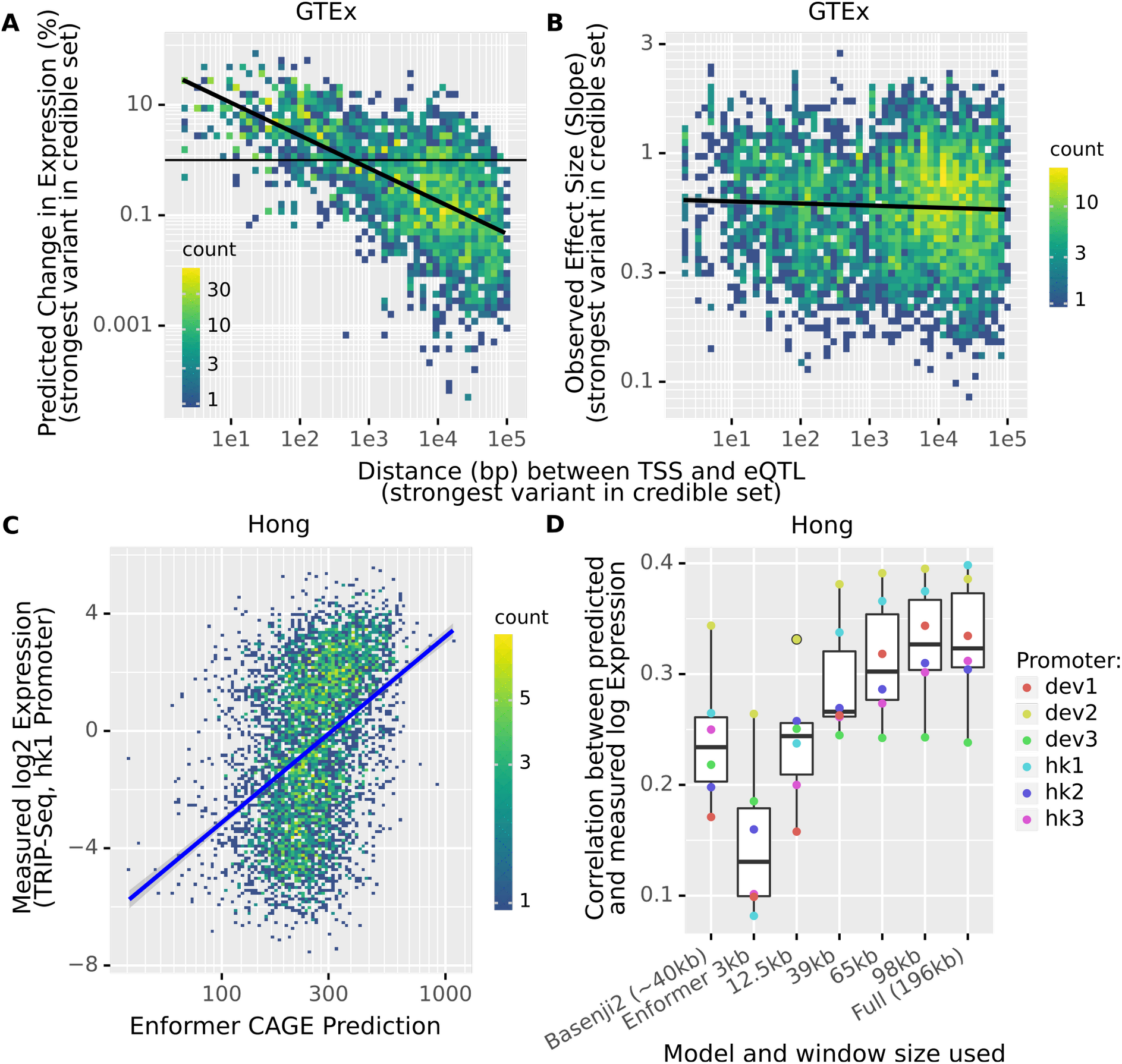
Enformer attributes no meaningful impact to distal eQTLs and performs poorly on tasks where long-range information is crucial. **A)** The measured and predicted changes in gene expression (expressed as an unsigned percentage) due to eQTL variants are plotted as a function of the distance between the (canonical) TSS and the eQTL. To account for linkage, we always take the maximal effect of all variants in the credible set. This predicted effect quickly decays with distance. **B)** The GTEx eQTL normalized effect size is plotted against the distance to the TSS. We observe no systematic decay with distance. **C)** Enformer struggles to predict the overall impact of the genomic environment on expression of the hk1 promoter, as measured by TRIP-seq. Note that this is the promoter for which the model performs best. **D)** Performance of Basenji2 and Enformer on the TRIP-Seq data. For Enformer, we computed predictions after restricting the input window, as previously. We find that most of the (limited) signal on this task once again derives from the proximal 20% of the input sequence.

### Current deep models mostly cannot predict the general impact of the genomic environment

To further probe the ability of Enformer to correctly account for distal sequence context in its predictions, we used TRIP (thousands of reporters integrated in parallel) sequencing data compiled by Hong et al. [27]. In this experiment, six different short promoter fragments were integrated at thousands of locations across the genome and then their activity at each location was measured. This assays the impact of the overall genomic environment on promoter activity. We replicated this experiment in-silico by inserting the fragments at the same locations and recording the predicted expression.

Consistent with the results described previously, Enformer perfectly ranked the promoters according to their median expression (Fig S15). However, its ability to predict the variation in expression of individual promoters when integrated at different genomic locations was quite limited (r = 0.24-0.4 depending on the promoter, Fig 6C,D, Fig S16), albeit statistically significant. Moreover, Enformer once again outperformed Basenji2 on this task, even when it was limited to the same receptive field (Fig 6D).

Hong et al. also tested the impact of placing 676 different promoters in 4 different locations in the genome (Patch-MPRA). When we reproduced this experiment in Enformer, we found very coherent results: (1) Enformer predicts the average impact of promoters well (r = 0.73, Fig S17), (2) the promoters alone explain more variation in Enformer predictions than in the experimental data (69% vs 55%), and (3) only intermediate-strength promoters, but not strong promoters, are predicted to be strongly affected by the genomic environment (Fig S18). Of note, this is somewhat consistent with the experimental data, but there the effect is far less pronounced (Fig S19).

Admittedly, the fact that both Basenji2 and more so Enformer have any predictive power on the TRIP-seq data at all is impressive since it requires predicting the impact of a rather extreme and unnatural modification of the genome. Nevertheless, the limited performance on this data further indicates that more research is needed to achieve models of transcriptional regulation which properly capture the effects of distal elements and genomic context.

### Enformer’s promoter-enhancer logic is (mostly) multiplicative

A remarkable result, obtained independently by Bergman et al. [12] and Hong et al. [27], is that promoters and enhancers generally act multiplicatively on transcriptional input. We asked whether Enformer qualitatively captured the same rule. To this end, we designed an in-silico experiment. This is not trivial to test because, as discussed previously, the promoter usually dominates all other sequence elements in Enformer. Therefore, we selected endogenous triplets of promoter, enhancer and sequence background which were such that the enhancer had a notable predicted impact on the promoter in that particular background (Methods). We then predicted for every combination of these promoters (n = 89) and enhancers (n = 115) in each background (n = 32). Remarkably, the Enformer predictions could be well approximated as the product of promoter strength, enhancer strength, and background strength (77% of the variance, Methods).

If we focus on individual backgrounds, we find that in some cases the enhancers can explain up to 25% of the variation in predicted expression at the promoter (Fig S20). Moreover, except for a few backgrounds where the variation in expression can be almost completely explained by a background strength, we find that the enhancer strengths are highly correlated across backgrounds (Fig S21). We find very similar results if we focus on each individual promoter (Fig S22). Thus, it appears that the background and promoter determine whether enhancers have any impact, but if they do, this impact is consistent.

Altogether, this analysis showed that Enformer qualitatively agrees with the recently reported multiplicative model of promoter and enhancer effects, despite quantitatively underestimating the effects of distant regulatory elements.

## Discussion

Here we have performed the most extensive benchmark of sequence-based models of gene expression to unseen data. Specifically, we compared the predictions of the models against two large-scale observational RNA-seq datasets of adult and developmental tissue, as well as five deep perturbation assays, comprising designed reporter assays and CRISPRi screens. With our approach, we specifically probed to what extent current sequence-based models can account for the regulatory role of promoters, enhancers, eQTLs, and genomic environment. This allowed us to evaluate the generalization power of these models. We found that current sequence-based models show a remarkable ability to predict the expression associated with human promoters. However, we observed that even Enformer, with its very wide receptive field, can account for distal regulators only to a limited extent.

We will now discuss in more detail our findings and provide suggestions on how the field could use and improve sequence-based models of gene expression.

### Do current sequence-based models learn causal effects?

Across a wide range of analyses, we repeatedly observed that current sequence-based models, particularly Enformer, predict the impact of different promoters very well. It does not matter if the strength of a promoter is measured in its proper genomic context, in a completely different context, or even in a plasmid -Enformer predictions of expression generally will show substantial correlations with these measurements.

Moreover, we observed that Enformer often correctly predicts the impact on expression of diverse promoter modifications. These include single-nucleotide variants, addition or removal of TF-binding sites, and in some cases mutagenesis or shifts of core-promoter motifs. This is strong evidence that the model captures causal determinants of the promoter.

In contrast to the strong performance for promoters, the model underperforms when it comes to accounting for the expression effect of enhancers, particularly if they are distal. However, we note that, since our analysis was focused on expression, we did not analyze whether the deep models can correctly account for the causal impact of enhancer variants or perturbations on epigenetic marks at the enhancer itself. Performing a similar analysis as we have done but focused on ATAC-Seq or CHIP-Seq-based MPRA or allele-specific binding data [28] could shed light on this question.

Accordingly, our results should not be construed to mean that such models do not further our understanding of enhancer biology or enhancer variants. We focused on showing that these models attribute far too little importance to enhancers when predicting gene expression. Indeed it has been repeatedly demonstrated in the literature that using the full range of epigenetic predictions can provide real added value in enhancer prioritization or variant fine-mapping [8–10,29].

### Do current sequence-based models learn long-range effects?

Overall, we found limited evidence that the sequence-based models we studied make use of long-range information. For Enformer, the model with the largest receptive field (196kb), we found that we can safely remove two-thirds of the input sequence, with minimal impact on predictions. As we saw from the distribution of eQTLs and validated enhancers this does to some extent reflect the underlying biology. Indeed, most strong regulatory elements do seem to be located relatively close to their target gene. However, the datasets we analyzed clearly showed that distal elements with large causal impacts on expression exist and that Enformer will generally strongly underestimate their impact. As a result, the majority of validated enhancers and eQTLs do not have a meaningful impact on Enformer predictions of gene expression.

We hypothesize that a large part of the problem is the escalating class imbalance: The ratio between actual regulatory elements among all candidate regulatory elements decreases with increasing distance from a gene. Perhaps, the model responds to this worsening signal-to-noise ratio by down weighting distal elements and thus, in some sense, distributing their effect. One possible underlying reason is the model being unable to correctly identify which distal regulatory elements will be functional in a certain cell type. Another non-exclusive, possibility is that the model is unable to correctly link distal regulatory elements to their target genes. In other words, is the source of difficulty the complexity of the enhancer code or is it the complexity of the “folding code”? Disentangling these issues will help to determine which additional training data and modeling assumptions are most likely to yield substantial improvements.

### Do current sequence-based models learn between-condition regulation?

We showed that Enformer, and to a lesser extent also Basenji2, predictions do exhibit significant correlations with measured between-condition differences in expression. Thus, clearly, the models capture some signal on this task. However, the between-condition predictions are less impressive, yet decent, than the between-gene ones.

We showed that this is partially because even Enformer has not yet reached the level of precision necessary to predict between-condition variation, which is generally of a smaller magnitude than between-gene variation. Thus, in principle, as models improve overall, they should particularly improve on between-condition tasks.

Our between-condition analysis was performed using endogenous gene expression and not using controlled perturbation experiments. For this reason, we cannot exclude the possibility that some of the between-condition predictive power is correlative. Furthermore, we cannot determine whether Enformer struggles at between-condition predictions precisely because of its limited understanding of distal regulatory elements. Since distal enhancers appear to mostly be a feature of multicellular organisms, one of their main roles could be to control cell-type specific expression. Deep perturbation assays performed across multiple cell types are needed to address these issues.

### Implications for in-silico investigations of gene expression

Our results indicate that sequence-based models can now be usefully employed to study promoter mutations in silico. This could prove useful for fine mapping in GWAS studies, for the detection of possibly pathogenic variants in rare disease diagnostics and oncology settings, and to study the evolution of human promoters. In-silico experimentation and interpretation techniques may even yield new candidate motifs or mechanistic hypotheses, which can then be tested in future designed MPRA studies. Assuming sufficient GPU resources, we recommend using Enformer for these tasks, as it clearly outperformed all other methods. However, some limitations of Enformer should be considered. Because it predicts in 128bp bins, Enformer cannot be used to predict the exact locations of TSS sites. Also, the model cannot reliably predict the directionality of a promoter because it is trained to predict similar values for both strands.

We found that variant effect predictions on gene expression from Enformer appear to be miscalibrated when comparing between promoters. Thus, genome-wide rankings of promoter variants made using Enformer may be misleading. We cannot fully exclude the possibility that this is an artifact of the plasmid-based measurements, rather than a failure of the model. Nevertheless, if a genome-wide ranking is the goal, using the predicted DNase variant effects (or a mixture of CAGE and DNase) appears to be the safer choice.

Our results indicate that predicting the impact of a distal variant on expression at the TSS rarely leads to meaningful results.

Finally, we note that, in addition to using sequence-based models to analyze variants or to prioritize enhancers, they also have potential use as an in-silico experimental platform to explore more abstract biological questions. A recent study, for example, used a deep model to analyze the impact of helical periodicity on TF binding [30]. As models mature, open questions in epigenomics, such as the pioneer factor hypothesis, the billboard vs. enhanceosome discussion, and others may become amenable to in-silico analysis. This could help to design more targeted experiments and also aid in the interpretation of experimental results.

### Implications of our study for model development

The recent trend in deep supervised models of regulatory genomics has been towards expanded receptive fields, through the use of dilated convolutions and attention. These expansions have usually been associated with increased performance, suggesting that the wider receptive field is the main cause of the improvement.

Paradoxically, however, we observed that Enformer, despite its huge receptive field, derives most of its predictive power from a small fraction of the sequence. Moreover, Enformer substantially outperformed Basenji2 even when it is restricted to the latter model’s input window and even on tasks where the receptive field size is irrelevant (such as plasmid-based saturation mutagenesis). Therefore, it is unclear to what extent the large receptive field of Enformer actually contributes meaningfully to its predictive power. Perhaps, similar to results in natural language processing [11,31], the mere amount of parameters combined with the transformer architecture, is the driving force behind the improvement.

How could a very good model of gene expression that does account for distal regulators be built? Given that Enformer already achieves close to replicate predictions on CAGE in ENCODE, it is likely that data complementary to ENCODE is required. Which data will help the most depends on the exact nature of the problem. If the issue is that Enformer does not correctly interpret the enhancer code, then adding more epigenetic, as well as expression, data from more cell types and more species should be most beneficial. More cell types provide more variation in enhancer activity, whereas more species introduce not just variation in activity but also variation in functional enhancer sequence. Recent advances have shown that, if done at a sufficient scale, perturbation experiments can also be used to train models [32–34], although further progress will require modeling advances to seamlessly integrate global and targeted assays.

If the main difficulty is folding, then adding the prediction of Hi-C data as an auxiliary task, or integrating a model which can predict Hi-C, would be the natural solution. Promising models predicting chromosomal contacts from sequence have been obtained [35–37], but integration with gene expression prediction models remains to be explored.

## Methods

### Models used

Basenji1/2 have a very similar structure as Enformer. Accordingly, Basenji1/2 predictions were always made in exactly the same way as Enformer predictions -the only difference being a smaller input window. We used the pretrained Basenji2 model from: https://storage.googleapis.com/basenji_barnyard/model_human.h5. For Basenji1, we used the model from the Kipoi model repository [38].

Xpresso only predicts a single value per sequence. Moreover, it is strand-specific and it expects the TSS to be at a particular position in the sequence. In cases where the TSS is known, we thus made only one prediction with Xpresso, namely with the TSS at the correct location and on the correct strand. In the Segal dataset, where the exact TSS is not known, we tried a number of offsets and selected the best ones. We used the Xpresso model from the Kipoi model repository, which does not account for RNA half-life covariates.

To get Expecto predictions of variant effects, we used the web interface at: https://hb.flatironinstitute.org/expecto/.

### Endogenous Expression: GTEx and Cardoso-Moreira et al

We collected gene expression measurements for GTEx tissues from the GTEx consortium webpage and for different development stages from ArrayExpress [39] (E-MTAB-6814). Note that this data was already normalized for sequencing depth and gene length. We log-transformed the data, adding a pseudo count of one. We exclude mitochondrial genes and the Y chromosome. We also exclude all genes which are never expressed in our data.

We extracted the reference genome sequence (hg38) around each gene, centered on the canonical TSS (i.e. the TSS of the ENSEMBL [17] Canonical transcript), and computed the predictions of expression at this TSS for each of the models.

Because Enformer/Basenji2 provide CAGE predictions only for certain ENCODE cell lines, which do not permit a 1:1 matching to GTEx tissues, we instead perform the matching using ridge regression. Specifically, we fit L2 regularized regressions for each tissue such that:

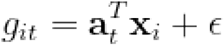

Where *g*_*it*_ is the (log-transformed) expression of gene i in tissue t, **a** _**t**_ is the vector of learned weights for tissue t and **x**_i_ is the vector of (log-transformed) Enformer/Basenji2 CAGE predictions for gene i. Note that the intercept is zero by construction (see below).

To fit these regressions, we split the genes into train and test-set, whereby every gene which is fully enclosed in an Enformer test-set region is included in the test set, and all genes which never intersect any test-set region go to the train set. Genes which intersect a test-set region, but are not contained by it, are excluded. In this way we prevent contamination of the held out test-set.

To make between-condition comparisons meaningful, we compute the mean for each tissue (or tissue-development-stage in the Cardoso-Moreira et al. [15] data) on the training set and remove this mean from both the training and the test set. This ensures that our regressions cannot learn tissue means, in a way that prevents leakage from the test set.

For Xpresso we follow the same procedure, but as this model is cell-type agnostic, the ridge regression only rescales the predictions.

We do not show the results for Basenji1 on this data, as Basenji1 used a different train-test split, thus the numbers are not comparable (nevertheless, it still performs worse than Enformer). Note that the same is true for Xpresso, but we consider it unlikely that this model overfits very much.

### Sequence context ablation study

We follow the same steps as we did previously for the GTEx and Cardoso-Moreira et al. data. The only difference being that we now generate predictions with different sequence window sizes around the TSS. We use the following window sizes: 1001, 3001, 12501, 34501, 39321, 49153, 65537, 98305 and 131073 bp. The last six correspond to a fifth, a fourth, a third, half and two-thirds of the total receptive field respectively. These windows are always centered on the TSS (so a window size of 1001bp means we extract the TSS site +/-500bp). As the windows are smaller than Enformer’s receptive field, we pad with “N” nucleotides on the flanks.

We train separate regression models for each window size, using the same train-test split as previously

### Class Imbalance

We downloaded ENCODE CREs [23] from https://screen.encodeproject.org/.

We used PyRanges [40] to define “Enformer-sized” (i.e. 196kb) windows around each gene of interest (i.e. the genes with validated enhancers or eQTL). We then intersected our sets of regulatory elements (eQTL, CRISPRi validated and non-validated enhancers and ENCODE CRE) with these windows and for each hit we recorded the distance to the gene. Note that we apply the same filters we applied to the eQTL data also to the ENCODE CRE, so as to make these sets comparable (these filters are discussed in the eQTL methods section). The validated and non-validated enhancers are comparable by design.

### Promoter determinants: Weingarten-Gabbay et al

We collected the data from the supplementary materials of the Weingarten-Gabbay et al. [24] manuscript and the construct sequences were kindly provided by the first author. We followed the general procedures outlined previously to construct sequences and compute predictions. As the exact TSS is, to our knowledge, unknown, we center predictions on the midpoint of the promoter fragments. As this is a K562 experiment, we use the K562 CAGE track as the predictor.

### Saturation Mutagenesis: Kircher et al

Vikram Agarwal kindly provided the Kircher et al. [13] data.

To generate a variant effect prediction with Enformer (and other models), we create two sequences: one centered on the reference allele and one on the alternative allele. We predict for both (averaged over strands, offsets, and the neighboring bins as always), and compute the log fold change in prediction centered on the variant (averaged over the neighboring bins).

Kircher et al. used a number of different cell lines in their experiment, depending on the locus. To match these cell lines to ENCODE tracks, we followed the same procedure as Avsec et al. [10]. Specifically, we used tracks (CAGE/DNASE) whose ENCODE descriptions contained substrings that correspond (more or less) to the cell line used for a particular locus:

- ‘HepG2’ for *F9, LDLR*, and *SORT1*
- ‘K562’ for *GP1BB, HBB, HBG1*, and *PKLR*
- ‘HEK293’ for *HNF4A, MSMB, TERT* and MYCrs6983267
- ‘pancreas’ for *ZFAND3*
- ‘glioblastoma’ for *TERT*
- ‘keratinocyte’ for *IRF6*
- ‘SK-MEL’ for *IRF4*

If there was more than one matching track, we averaged predictions over the matching tracks. This is different from Avsec et al., who instead extract principal components, but ultimately this procedure yields very similar correlations.

In cases where there was no match at all, we averaged predictions over all tracks.

### Promoter x Enhancer: Bergman et al

We collected the data and plasmid sequence from the supplementary materials of the Bergman et al. [12] manuscript. We employ the same filtering strategy, keeping only data points supported by at least 25 plasmids and at least 2 barcodes.

As this assay was conducted using a plasmid and has no clear analog in the endogenous genome, we reproduced the plasmid sequences in-silico and added N-padding on the flanks to adapt it to the Enformer input length. We placed the promoter and enhancer fragments into their respective locations in the plasmid and centered predictions on the midpoint of the promoter fragment, as the exact TSS is -to our knowledge -unknown.

Bergman et al. use their data to impute the intrinsic strengths of the promoter and enhancer sequences. For this, they fit a Poisson model with promoter and enhancer indicators:

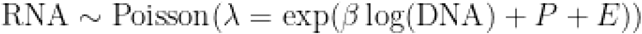

where RNA is the measured RNA count, DNA is the amount of plasmid used, *β* is a learned weight, and P and E are the promoter and enhancer indicators (“strengths”) respectively. We reproduced this analysis using the package statsmodels [41]. Note that we also fitted a log-linear OLS to this data, which gave very similar results, but we report the results of the Poisson model to stay faithful to the source material.

To impute the *predicted* promoter and enhancer strengths, we use a similar strategy. However, we do not fit a Poisson model, as Enformer predictions are not integer-valued. Instead we fit a Gamma model:

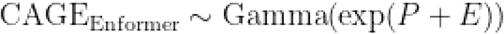

where CAGE is the Enformer CAGE prediction (for K562).

The Gamma distribution is often used to model the prior distribution of a Poisson lambda and arguably the Enformer prediction in natural scale is a Poisson lambda, as Enformer is trained using a Poisson loss function.

To ensure that our results are robust, we additionally fitted a linear regression to the log-transformed Enformer prediction:

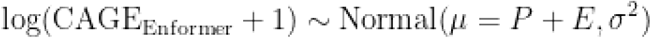

This gave very similar results.

Note that our calculations of explained variance refer to the variation in log expression. This is why they slightly differ from the ones reported in Bergman et al.

### Enhancer knockdown

We collected the data from the supplements of the respective manuscripts. Additionally, the sequences used for this benchmark in the Enformer paper were kindly provided by Ziga Avsec. For each gene, Avsec et al. [10] determined the TSS site with the highest predicted expression in K562. They then extracted sequences centered on these TSS, unless the distance between the gene and the enhancer is such that this was not possible given the receptive field size, in which case the TSS is shifted to accommodate the tested enhancers. Predictions are made at the TSS sites, as per usual.

For each reference sequence (with the enhancer intact) we create 25 “knockout” sequences, where we shuffle 2000 bp centered on the enhancer. In this way, we destroy the enhancer without changing the nucleotide composition of the underlying sequence. The predicted effect of enhancer knockout is then given by the average (over shuffles) change in predicted gene expression at the TSS.

### eQTL

We downloaded SuSie credible sets for GTEx eQTL from the EBI eQTL catalog [26]. We additionally downloaded GTEx eQTL normalized effect sizes from the GTEx portal [14,16].

We apply the following filters:

- We only consider protein-coding eGenes.
- We exclude credible sets which span more than 5kb
- We demand that each variant in the credible set can be scored by Enformer when the canonical TSS is placed in the center of the sequence. In other words, the furthest variant of a credible set must be no further than 98kb from the canonical TSS of the eGene, otherwise the entire set is excluded.
- We demand that all variants in the credible set are upstream of the canonical TSS. This is to exclude post-transcriptionally acting variants (i.e. NMD variants, splice variants, etc). If a credible set contains even one downstream variant, we exclude it.
- We exclude all eGenes which have a GENCODE annotated protein-coding transcript upstream of the canonical one. In this way, we ensure that the canonical TSS is always the closest (protein-coding) TSS of the eGene to the variant.

6141 credible sets pass our filters, thus providing a total of 14139 variants to test.

For each variant in each credible set, we then compute predictions for all CAGE tracks at the canonical TSS of the eGene (using our usual strategy of summing over neighboring bins and averaging over strands and small offsets). We then use the previously fitted ridge regressions to match these CAGE predictions to GTEx tissues. Next, we compute, for each variant, the change in prediction vis-a-vis the predictions made with the reference sequence. We define the eQTL effect as the effect of the strongest variant (i.e. the one leading to the biggest change in predicted expression at the canonical TSS of the eGene in the tissue of interest) in the credible set. This follows the usual assumption in the literature that generally only one variant in a linkage block will be causal. Our strategy fails if the eQTL effect is the result of an epistatic interaction between several variants in linkage -however testing this possibility would require testing all combinations of variants in a credible set, which would be prohibitively expensive. Moreover, we would expect that such cases are rare anyways.

### Trip-Seq (Hong et al.)

We collected the data and relevant construct sequences from the supplements of the Hong et al. [27] manuscript. We followed the standard procedure to replicate this experiment.

To compute the explained variances, we fit log-linear OLS models, similarly as we did for the Bergman et al. data.

### In-silico Multiplicativity Assay

To identify triplets of promoter, enhancer, and background where Enformer attributes importance to the enhancer in determining expression at the promoter, we returned to our in-silico reproduction of the CRISPRi data. We first selected promoter-enhancer pairs where:

- the enhancer is further than 3kb from the TSS
- the enhancer is within 90kb of the TSS
- the in-silico enhancer knockdown has a predicted impact of at least 30% (note that this includes both repressive and activating enhancers)

To cover a slightly bigger range of possible enhancer effects, we identified for the same genes some weak (< 1% predicted effect, despite being located within 20kb of the TSS) and intermediate strength enhancers (between 4% and 8%) predicted effect. We finally selected a few “non-functional” promoter-enhancer pairs where the enhancer had no real effect at all (< 0.1%).

This procedure yielded 89 unique promoters and 115 unique enhancers. To extract the promoter sequences, as exact boundaries are unknown, we take a 1kb window around the TSS. For the enhancer, we take a 2kb window around the enhancer midpoint.

Each promoter-enhancer pair comes with an endogenous sequence background, which also determines the distance of the enhancer to the promoter. Ideally, we would have tested every promoter-enhancer combination in every background, but the combinatorial explosion makes this computationally expensive - specifically because we have to predict for each sequence six times, i.e. for both strands and with small shifts, to limit the noise in the prediction. Thus we selected 32 backgrounds. We took 6 backgrounds from our set of non- functional triplets (to see if other enhancer-promoter pairs could be linked in such a background) and we selected 6 backgrounds where the enhancer location is far from the TSS location. The remaining backgrounds were selected from the strong triplets.

We then proceed as follows: for each background, we predict expression in K562 at the TSS for every combination of promoter and enhancer. Hereby, we always replace the endogenous promoter of this background with the promoter of interest and we replace the endogenous enhancer with the enhancer of interest. Thus, for a given background, the distance between promoter and enhancer is constant. We focus on K562 as most of the experiments on this topic were also performed in this cell type.

We get a total of 327,520 combinations (promoter x enhancer x background). We then fit log-linear models to explain the variation in log expression in this data. We first do this for the entire dataset, using an indicator variable for each promoter, each enhancer, and each background (236 parameters). This model thus assumes that the log expression of a certain promoter-enhancer pair in a certain background is determined by the innate strength of the promoter which is scaled by the background and the enhancer.

We next fit log-linear models to explain the expression variation for each background, across promoters and enhancers (32 backgrounds, with 10235 observations for each one). We use promoter and enhancer indicators (204 parameters). We also fit log-linear models to explain the expression variation for each promoter, across backgrounds and enhancers (89 promoters, with 3680 observations for each one). In this case we use background and enhancer indicators (147 parameters). Lastly, we correlate the enhancer parameters we estimated across promoters and across backgrounds.

## Declarations

### Ethics approval and consent to participate

Not applicable.

### Consent for publication

Not applicable.

### Availability of data and materials

The datasets analyzed during the current study are available in the Zenodo repository, https://doi.org/10.5281/zenodo.7076228. All scripts used in the analysis can be found under https://github.com/Karollus/SequenceModelBenchmark.

### Competing Interests

The authors declare that they have no competing interests.

### Funding

This study was funded by the German Bundesministerium für Bildung und Forschung (BMBF) through the Model Exchange for Regulatory Genomics project MERGE (031L0174A to JG and TM). The funders had no role in the study design, data collection, analysis, decision to publish, or preparation of the manuscript.

### Author contributions

AK and JG conceived the study and wrote the manuscript. AK analyzed the data. TM and AK designed, implemented, and tested the computational pipeline. All authors read and approved the final manuscript.

## Acknowledgments

We thank the authors of all the models we analyzed and all the assays we benchmarked against for making their code and data easily available. This study would not have been possible without their hard work. We specifically thank Shira Weingarten-Gabbay, Ziga Avsec, Vikram Agarwal, and David Kelley for providing advice on how to interpret their data and models.

## Notes

### Competing Interest Statement

The authors have declared no competing interest.

