## Supplementary Figures for "Current sequence-based models capture gene expression determinants in promoters but mostly ignore distal enhancers"

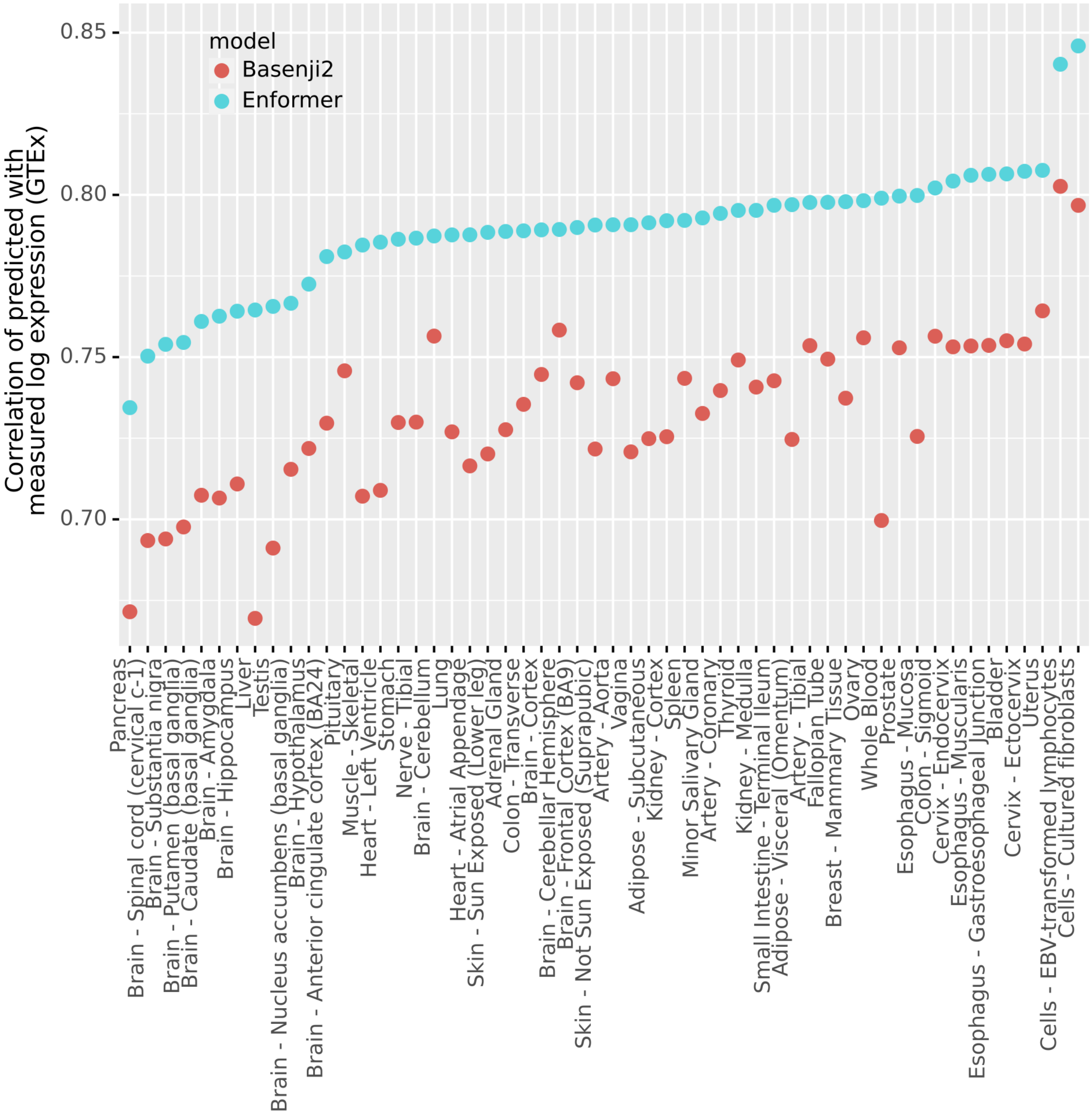


***Supplementary Figure 1****: Pearson correlation of predicted (using Enformer and Basenji2) and measured log expression for different GTEx [1] tissues. We see that both models perform best for cultured cell-lines, likely as these most closely resemble the ENCODE data and also are less complex than human tissues. Performance is lowest for Pancreas, most (but not all) brain tissues, liver and testis. In any case, Enformer always outperforms Basenji2.*

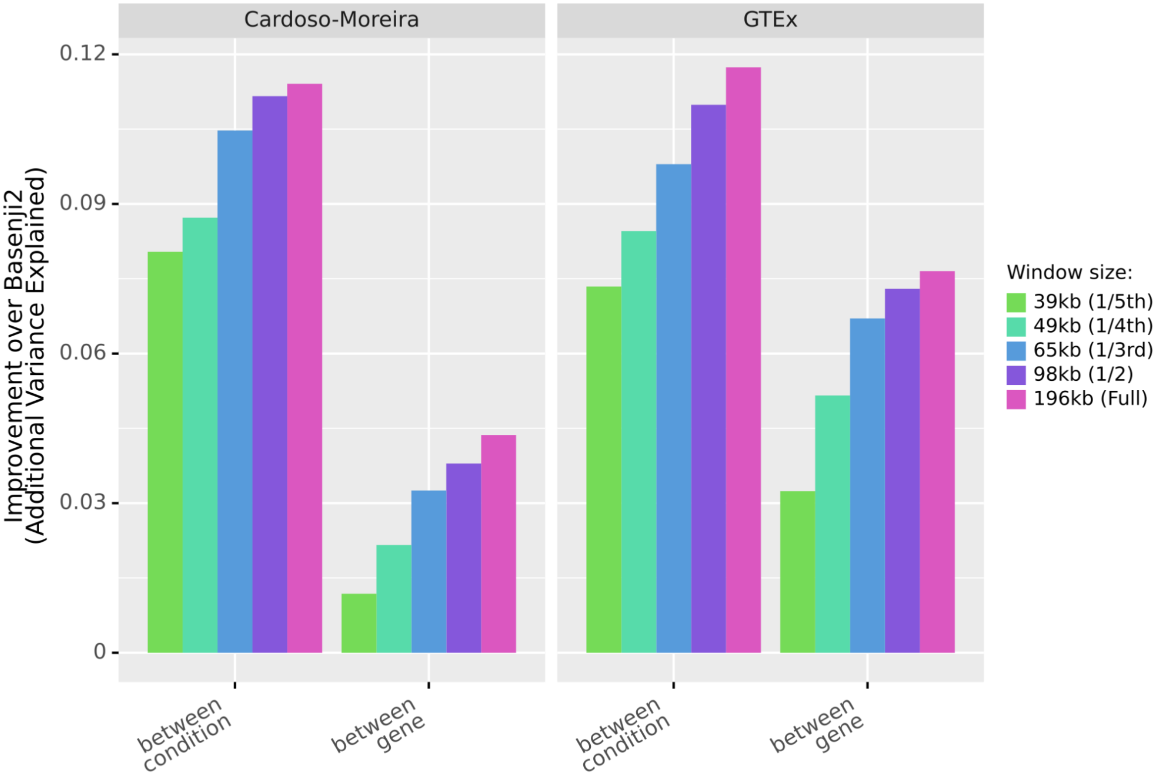


***Supplementary Figure 2****: the additional fraction of variance in log expression, both between-conditions and between-genes, explained by Enformer but not by Basenji2. We vary the amount of sequence context Enformer can use in its predictions, from 39kb (similar to Basenji2's receptive field of ~40kb) to the full 196kb. We see that more than half of the additional between-condition variance explained by Enformer is already achieved even when Enformer can only access the same sequence context as Basenji2. For between-gene, 30-40% of the improvement is achieved with the same sequence. The next 25kb of sequence (i.e. expanding the Basenji2 receptive field by ⅝) provides most of the remaining improvement. The remaining ⅔ of the Enformer receptive field (which represents an expansion of the Basenji2 receptive field of about 3-fold) contributes only a small additional improvement.*

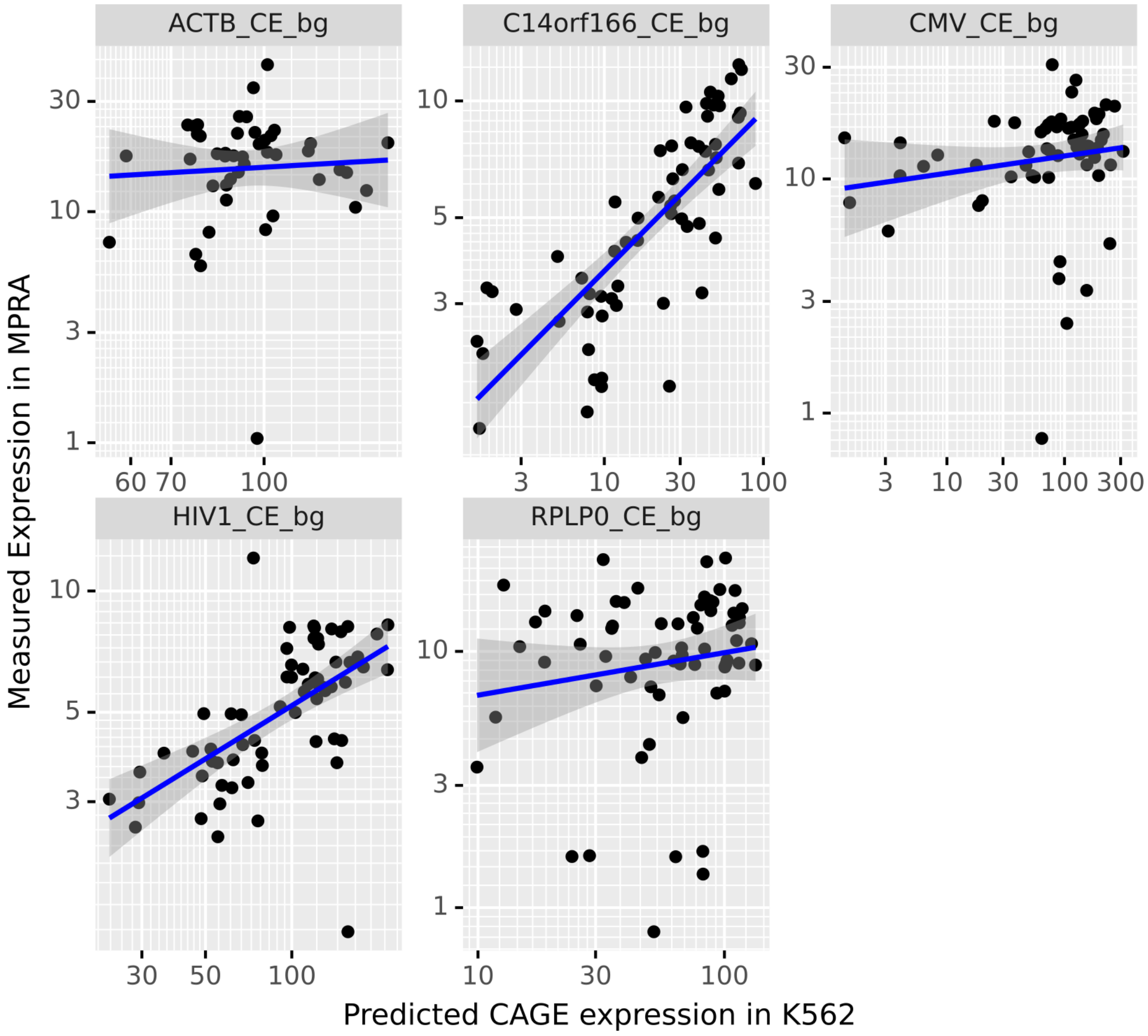


***Supplementary Figure 3****: Predicted vs Measured expression for an experiment where Weingarten-Gabbay et al. [2] inserted different combinations of core-promoter motifs (TATA, initiatior, BREu, BREd, MTE and DPE) five different backgrounds.*

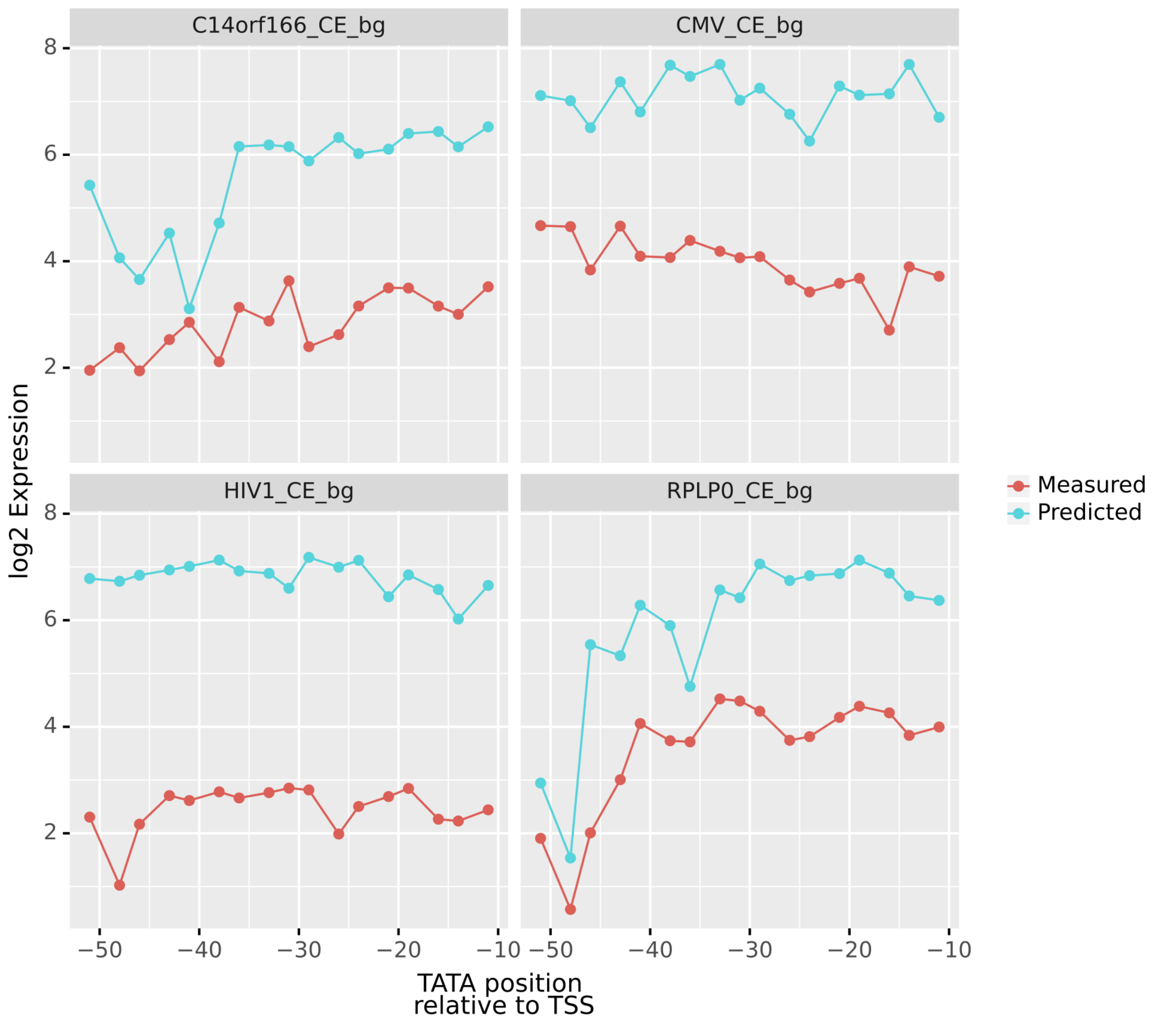


***Supplementary Figure 4****: Predicted vs Measured log expression for an experiment where Weingarten-Gabbay et al. [2] shifted the position of a TATA motif in four different backgrounds.*

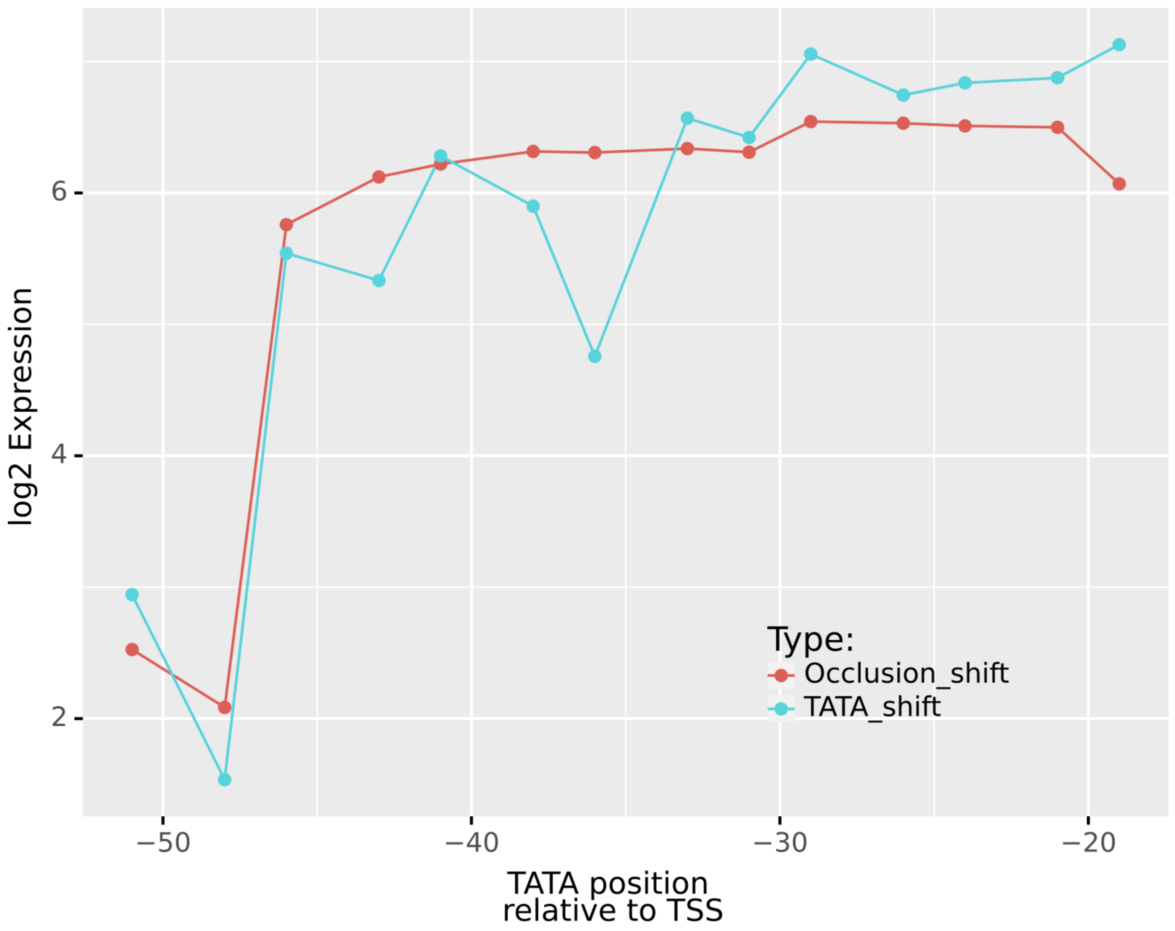


***Supplementary Figure 5**: Predicted log expression for an in-silico experiment where (blue) we shift a TATA box through the sequence of the RPLP0 promoter and (red) we shift a neutral motif (NNNNNNNN), which simply occludes the local sequence but adds no information, through the sequence. We see that the occlusion has very little impact, except around position -50, where it has almost the same impact as the TATA shift. This suggests that the effect there is not due to the TATA, but rather due to the local sequence that is perturbed if we place a TATA there.*

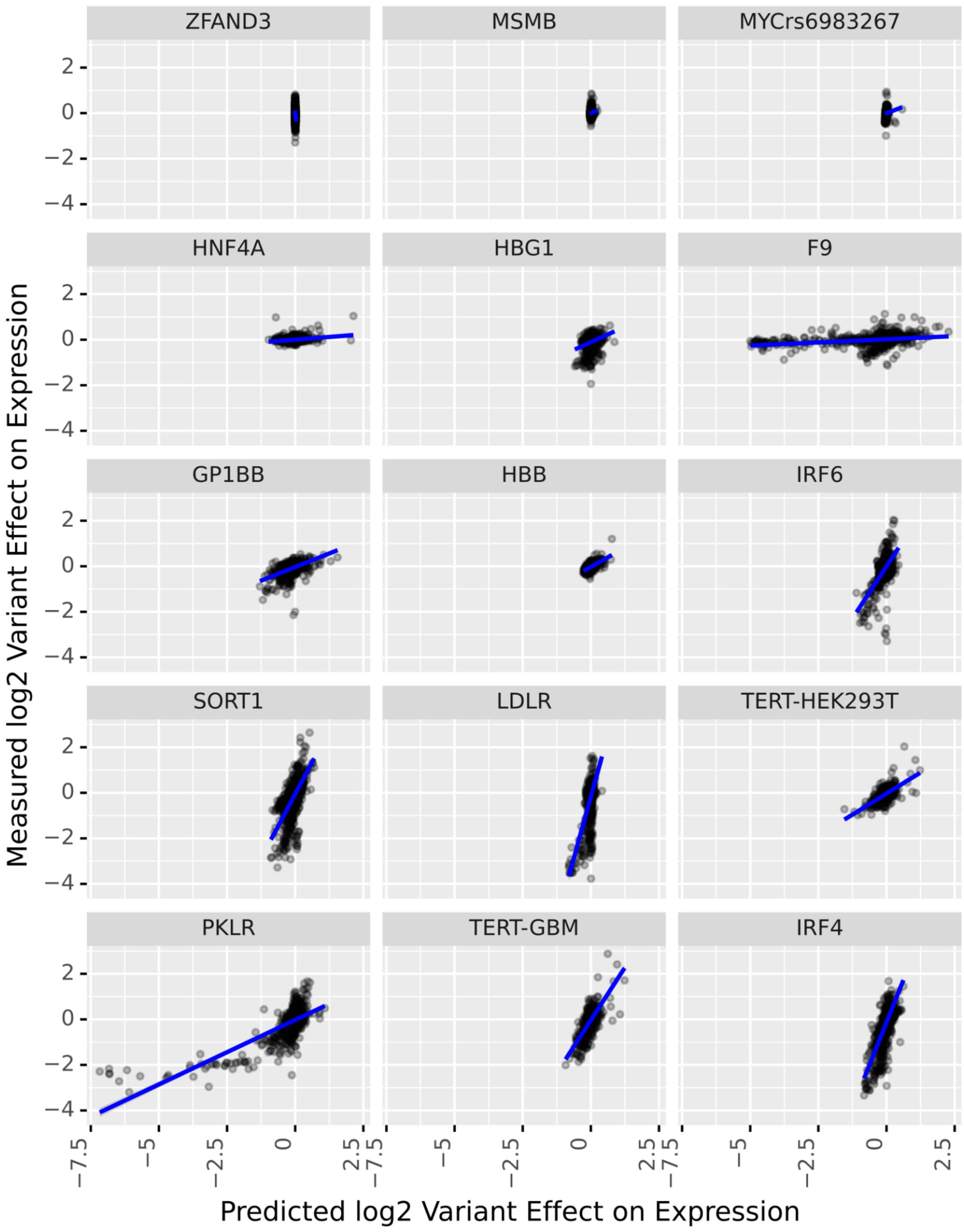


***Supplementary Figure 6****: Predicted vs Measured log2 variant effects for the Kircher et al. [3] saturation mutagenesis data, ordered (ascending) by the Pearson correlation. We observe that the best fit lines do not exhibit consistent slopes, suggesting a miscalibration between loci.*

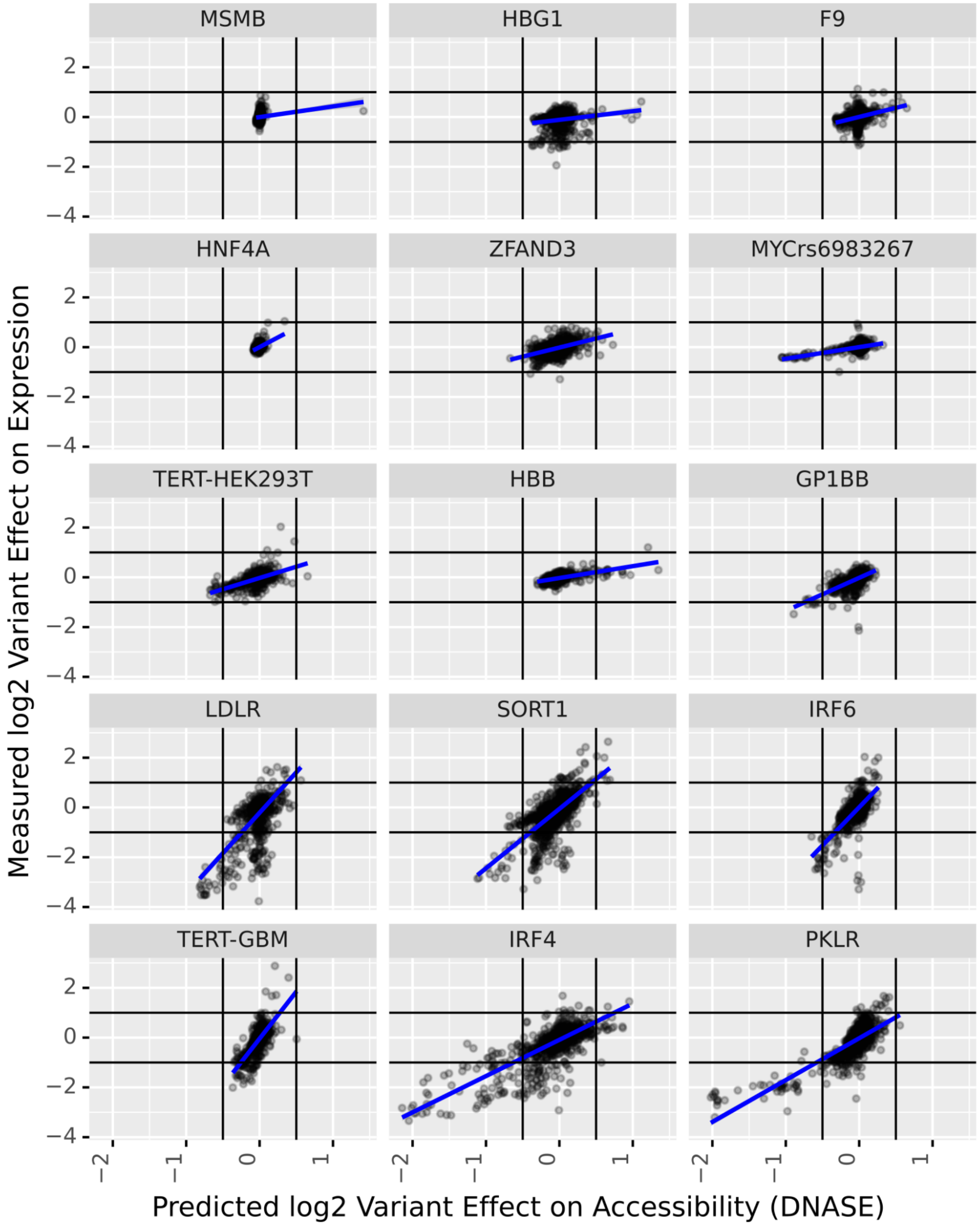


***Supplementary Figure 7****: Predicted vs Measured log2 variant effects for the Kircher et al. [3] saturation mutagenesis data, ordered (ascending) by the Pearson correlation. Note that here we use DNASE-predictions to compute the predicted variant effects. The slopes are more consistent. Note, however, that the effects are mostly too small: a predicted log2 fold change of ~0.65 usually corresponds to a measured log2 fold change of ~1.*

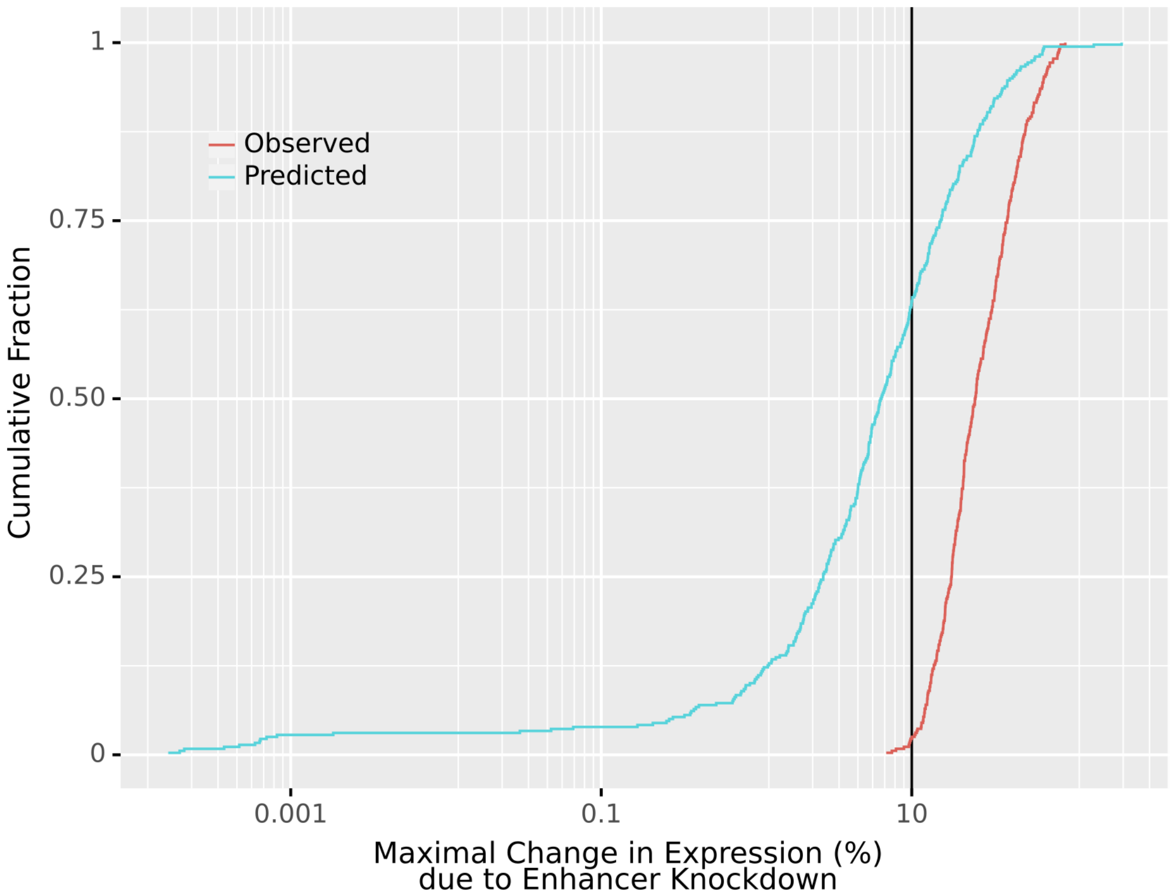


***Supplementary Figure 8****: Distribution of the maximal change in (predicted/measured) expression (%) induced by any of the enhancers tested by Fulco et al. [4] or Gasperini et al. [5] for genes which had at least one validated enhancer. We observe that for ~60% of genes, no tested enhancer (whether validated or not) had a predicted effect in excess of 10%.*

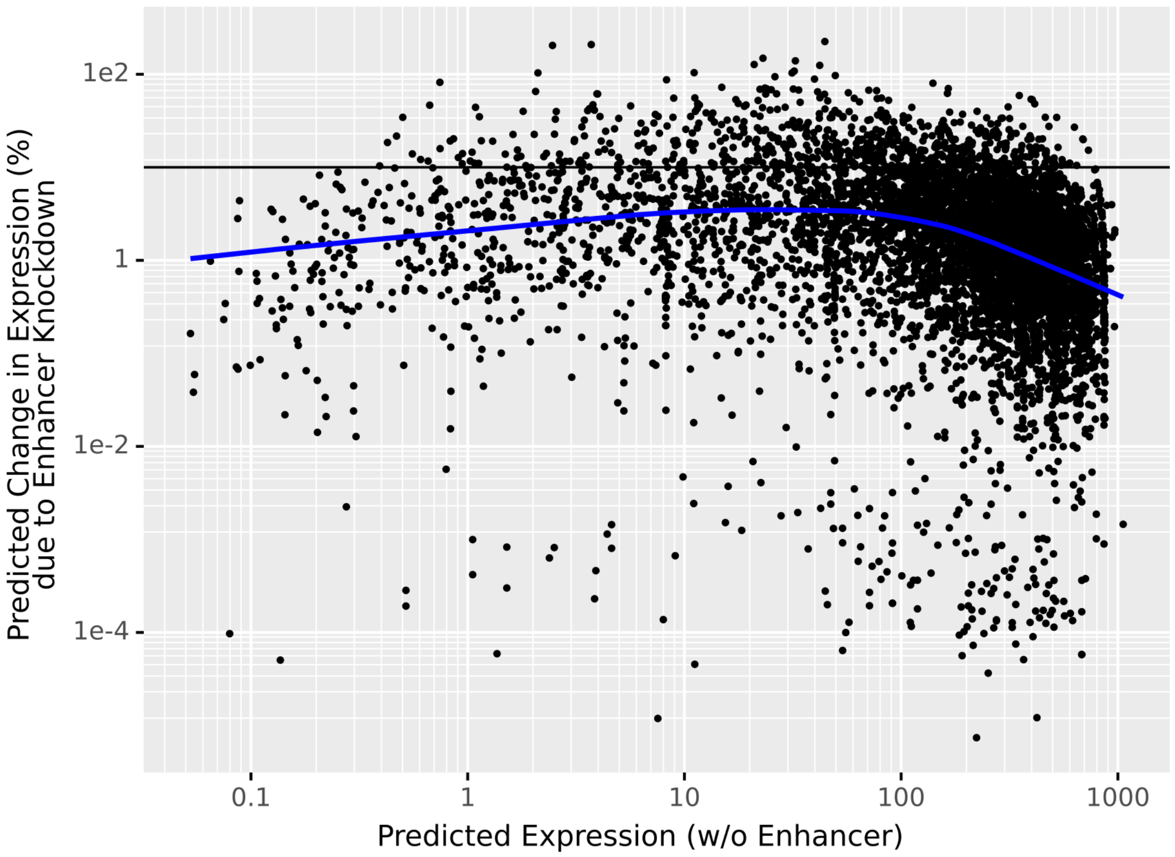


***Supplementary Figure 9****: The predicted (relative) impact of enhancer knockdown vs. the predicted expression at the TSS after enhancer knockdown. We see that genes with higher predicted basal expression (without a given enhancer) also generally react less to this enhancer.*

OLS Regression Results

| | | | |
|---|---|---|---|
| **Dep. Variable:** | log_pct | **R-squared:** | 0.249 |
| **Model:** | OLS | **Adj. R-squared:** | 0.248 |
| **Method:** | Least Squares | **F-statistic:** | 704.8 |
| **Date:** | Sat, 03 Sep 2022 | **Prob (F-statistic):** | 0.00 |
| **Time:** | 17:28:35 | **Log-Likelihood:** | -7007.6 |
| **No. Observations:** | 6394 | **AIC:** | 1.402e+04 |
| **Df Residuals:** | 6390 | **BIC:** | 1.405e+04 |
| **Df Model:** | 3 | | |
| **Covariance Type:** | nonrobust | | |

| | **coef** | **std err** | **t** | **P>\|t\|** | **[0.025** | **0.975]** |
|---|---|---|---|---|---|---|
| **Intercept** | 4.4606 | 0.103 | 43.297 | 0.000 | 4.259 | 4.663 |
| **validated[T.True]** | 0.1093 | 0.033 | 3.309 | 0.001 | 0.045 | 0.174 |
| **log_dist** | -0.8633 | 0.022 | -39.690 | 0.000 | -0.906 | -0.821 |
| **log_prom** | -0.2561 | 0.013 | -19.362 | 0.000 | -0.282 | -0.230 |

| | | | |
|---|---|---|---|
| **Omnibus:** | 1576.225 | **Durbin-Watson:** | 1.531 |
| **Prob(Omnibus):** | 0.000 | **Jarque-Bera (JB):** | 4067.675 |
| **Skew:** | -1.332 | **Prob(JB):** | 0.00 |
| **Kurtosis:** | 5.858 | **Cond. No.** | 58.3 |

***Supplementary Figure 10****: Results of a linear model which attempts to explain the predicted log-percentage change due to enhancer knockdown as a function of the log distance (log_dist), log of the basal gene expression (log_prom) and a indicator denoting whether a given tested enhancer was CRISPRi validated. Validated enhancers tend to have slightly higher predicted effects when controlling for distance and basal expression.*

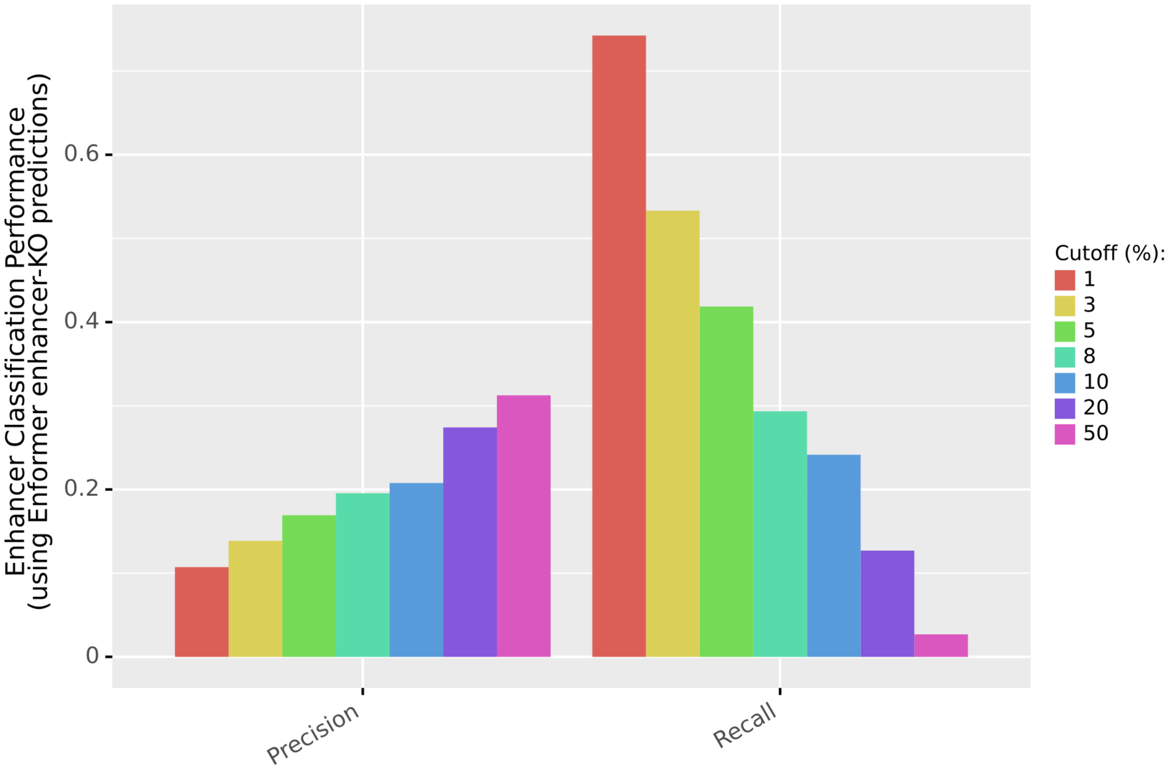


***Supplementary Figure 11****: Enformer precision and recall for prioritizing CRISPRi validated enhancers for different cut-offs. To achieve more than 50% recall, we need to classify as positive every enhancer with 3% or more predicted impact. Even with a 1% cutoff, we still miss almost 30% of validated enhancers.*

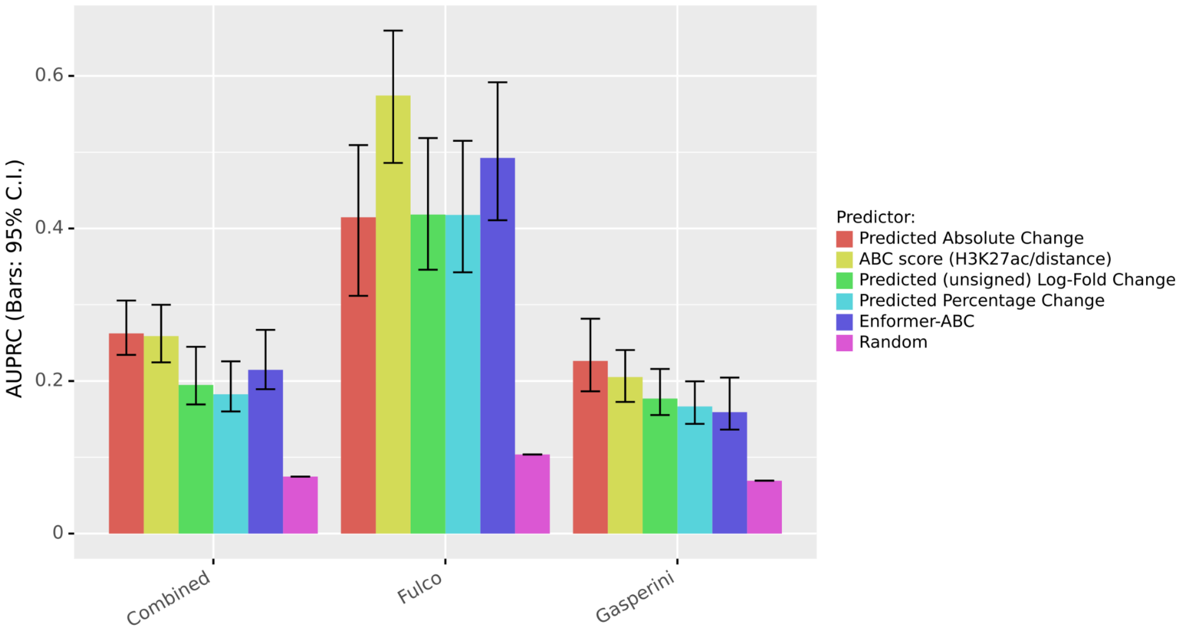


***Supplementary Figure 12****: Area under the precision recall curve for different methods of prioritizing CRISPRi validated enhancers. These include using the Enformer predicted absolute change in expression upon enhancer knockdown (red), the predicted relative change in expression (as percentage, green, or unsigned log-fold change, turquoise), a version of the ABC score [4] which does not use Hi-C measurements (see Avsec et al. [6] supplements for a discussion of this metric), the Enformer-ABC score (see below) and the random performance (i.e. the class balance). For the Gasperini data in particular, the absolute predicted change outperforms the relative predicted change, but this is likely an artifact.*

*The Enformer-ABC score is an analogue of the ABC score, but using Enformer DNASE and H3K27ac predictions centered on the enhancer location. In other words, the Enformer-ABC score of an enhancer is simply the geometric mean of predicted DNASE and HK327ac predictions, divided by distance. This Enformer-ABC correlates well (r = 0.7) with the non Hi-C ABC score used above. It performs slightly worse at prioritization than using the in-silico knockdown-effects, but it has the advantage that it seamlessly generalizes beyond the Enformer receptive field. It also performs worse at prioritization than the non Hi-C ABC-score, but it has the advantage that it may be responsive to enhancer variants without requiring new data. However, a thorough evaluation and hyperparameter optimization of this Enformer-ABC score is beyond the scope of this study.*

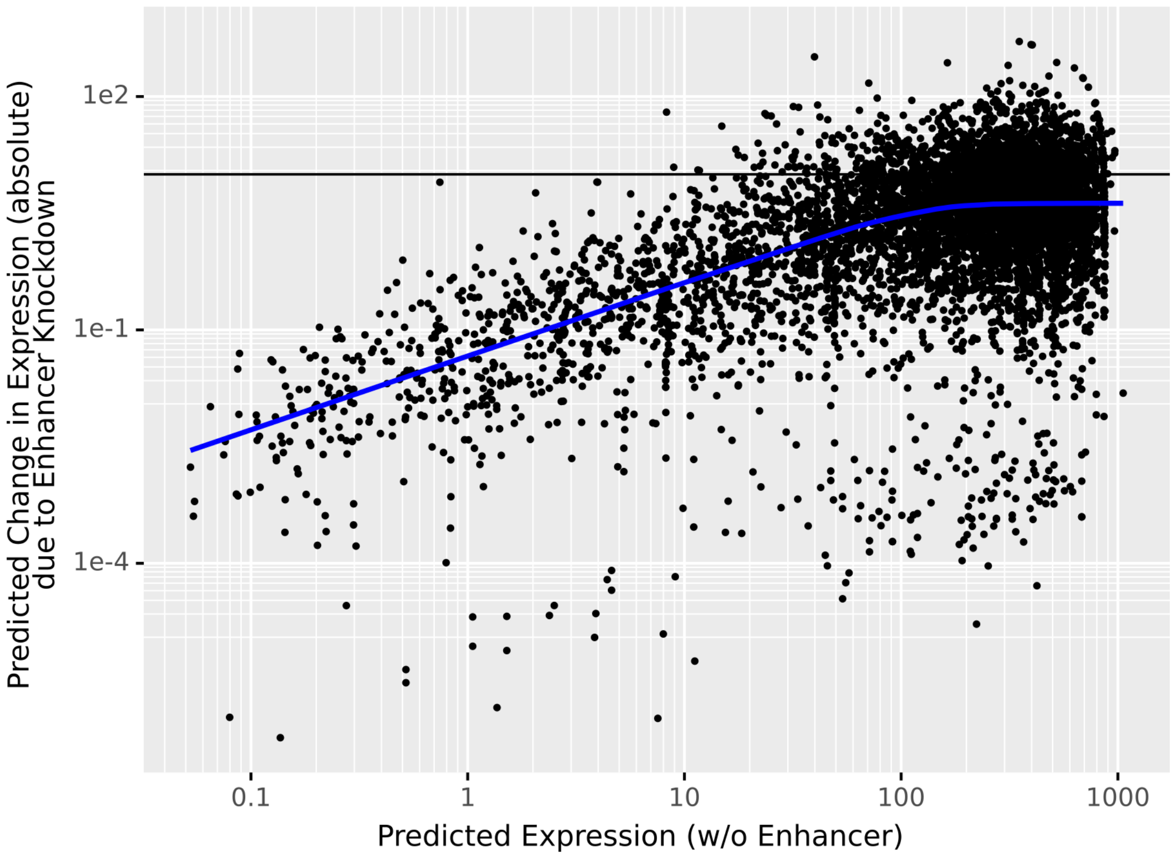


***Supplementary Figure 13****: The predicted (absolute) impact of enhancer knockdown vs. the predicted expression at the TSS after enhancer knockdown. We see that, up to a saturation point, the absolute effect rises mechanically as a function of the basal expression level.*

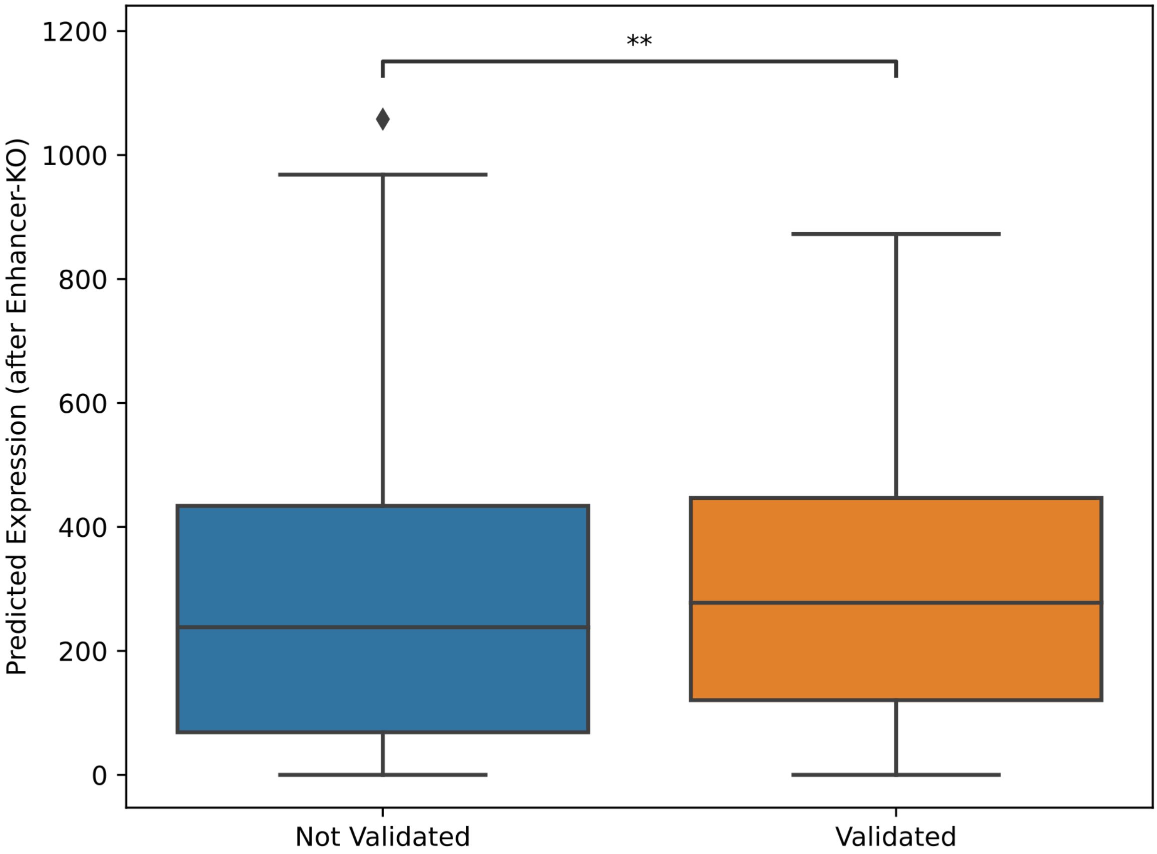


***Supplementary Figure 14****: We observe a small, but statistically significant (Wilcoxon test) association between the predicted basal expression of a gene and whether or not a given enhancer associated to this gene is validated. In other words, genes with higher predicted expression have more validated enhancers. This is somewhat unsurprising: the predicted expression correlates well with actual expression and perturbation assays generally have higher power to detect changes for more highly expressed genes.*

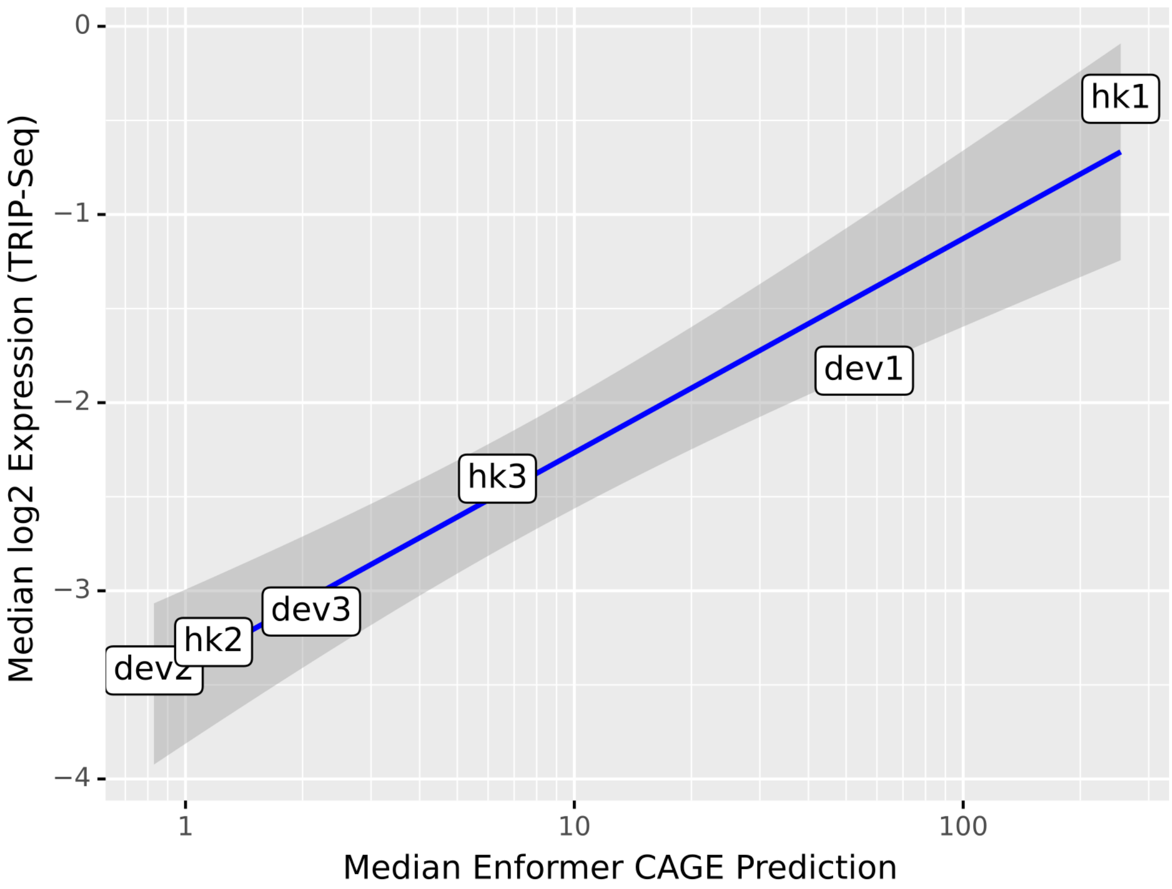


***Supplementary Figure 15****: The median, across genomic backgrounds, measured vs. predicted expression of each promoter tested in the Hong et al. [7] Trip-Seq experiment.*

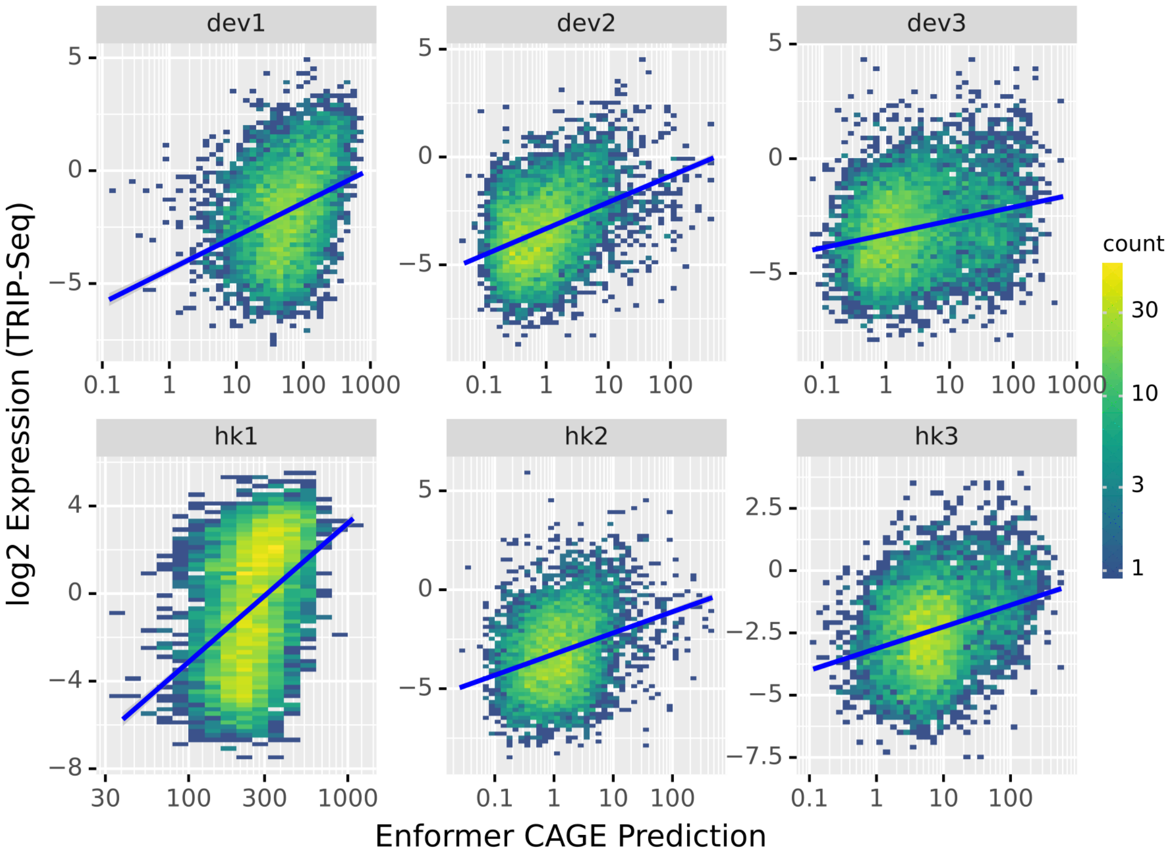


***Supplementary Figure 16****: The measured vs. predicted expression of each promoter at each location tested in the Hong et al. [7] Trip-Seq experiment. The correlation between Enformer predictions and observations tends to be poor.*

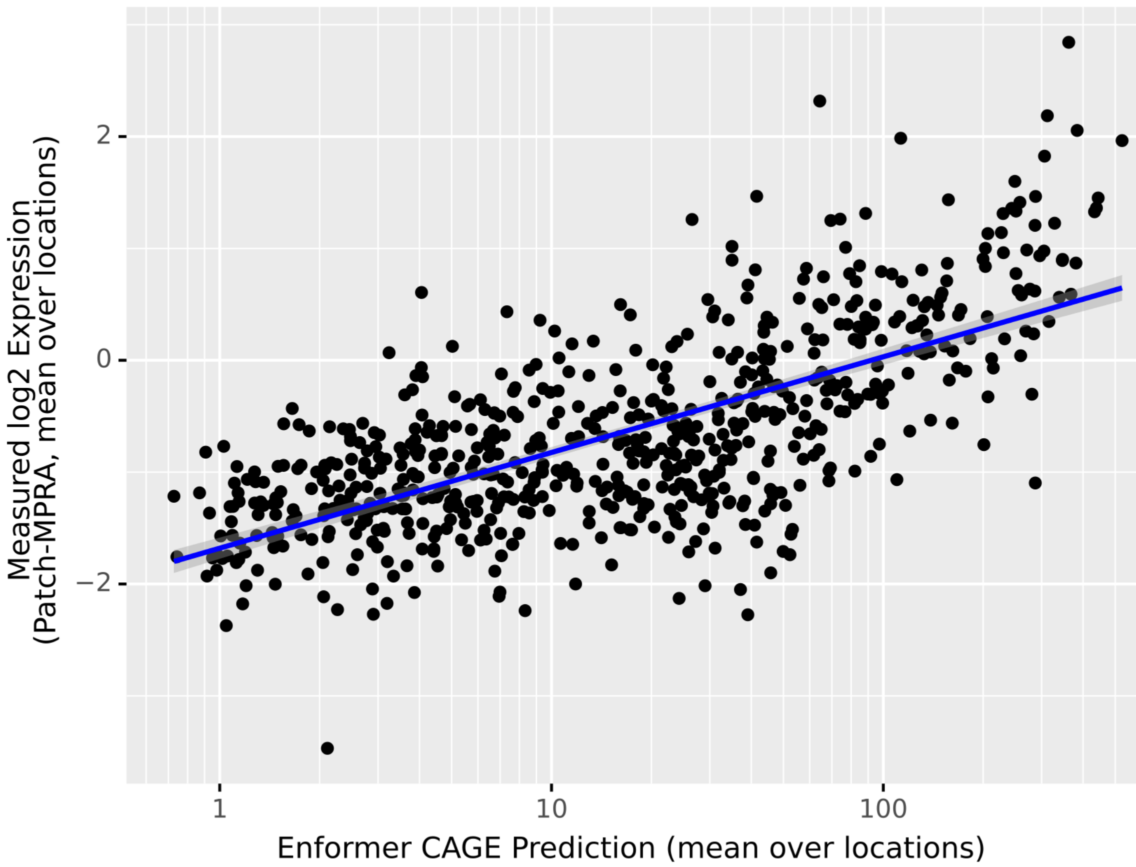


***Supplementary Figure 17****: The average, across genomic backgrounds, measured vs. predicted expression of each promoter tested in the Hong et al. [7] Patch-MPRA experiment. Once again, Enformer predictions of innate promoter strength match the measurements very well.*

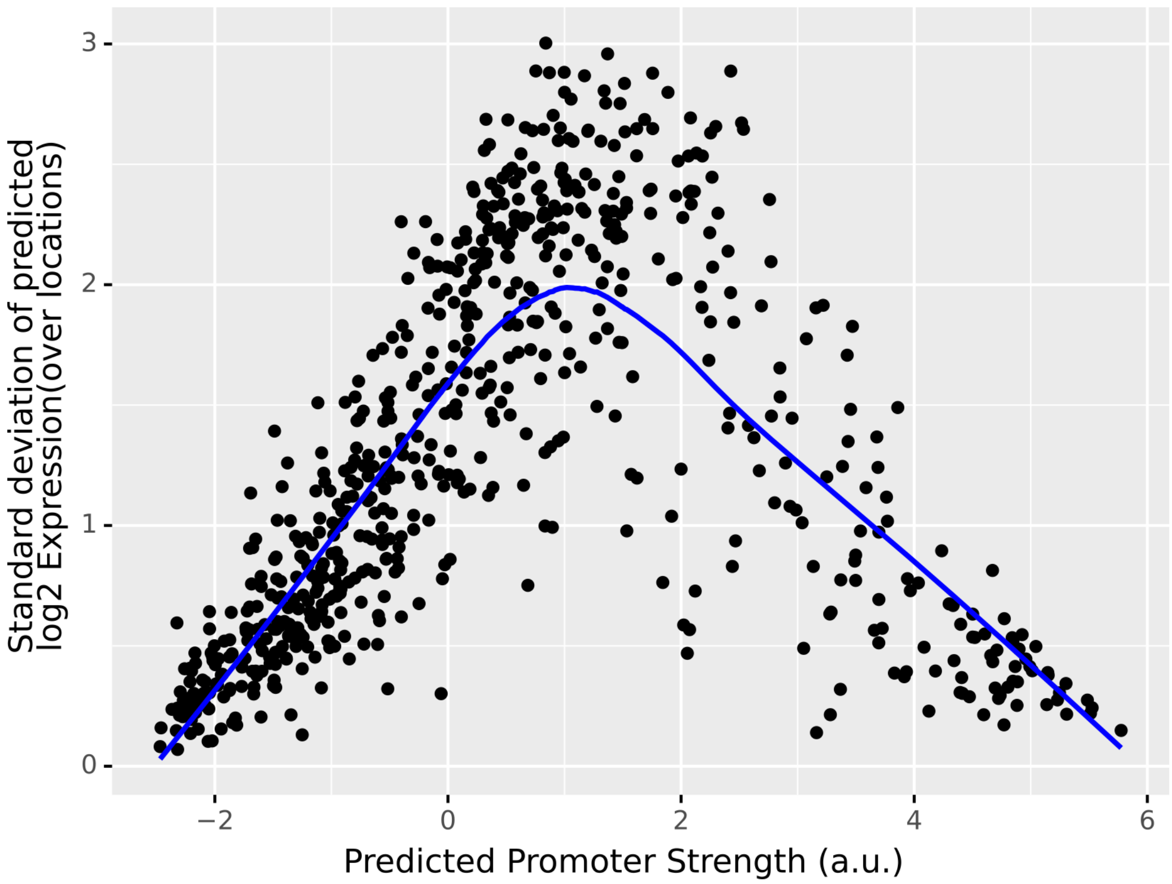


***Supplementary Figure 18****: The standard deviation of predicted expression for each promoter across backgrounds tested in the Hong et al. [7] Patch-MPRA experiment versus the predicted innate strength of each promoter (imputed with a log-linear model). We see that promoters Enformer considers strong overall react very little to the genomic background.*

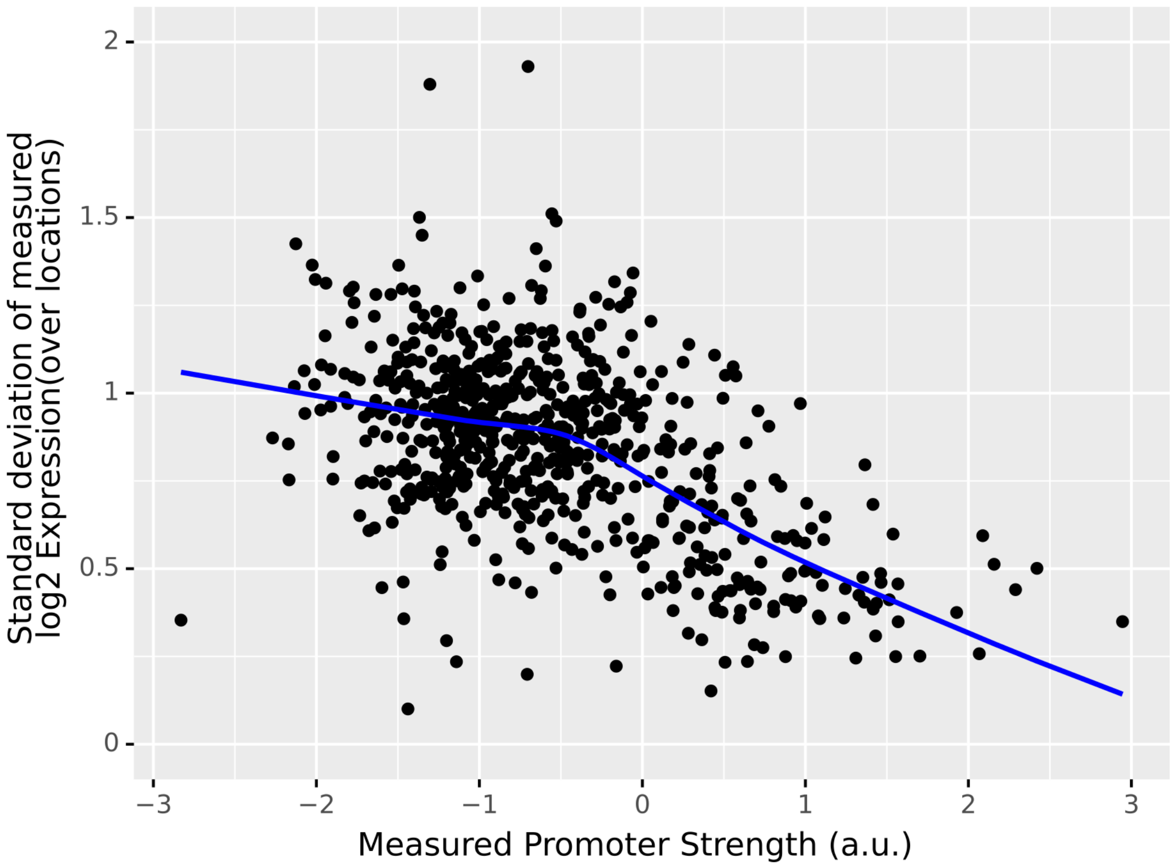


***Supplementary Figure 19****: The standard deviation of measured expression for each promoter across backgrounds tested in the Hong et al. [7] Patch-MPRA experiment versus the measured innate strength of each promoter (imputed with a log-linear model). Compare to supplementary figure 19. We see that in the experiment, stronger promoters also appear to react somewhat less to the background.*

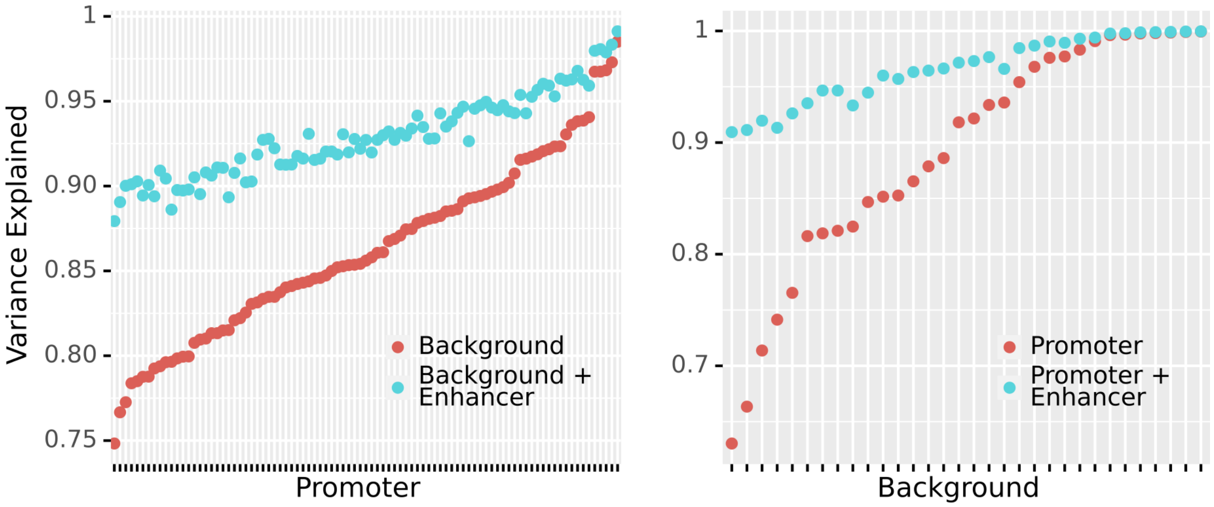


***Supplementary Figure 20****: The variance explained by a log-linear model for each promoter (using a background indicator or background and enhancer indicators as parameters) and for each background (using either only promoter indicators or promoter and enhancer indicators). We see that the amount of variance explained by the enhancer depends heavily on the promoter and the background.*

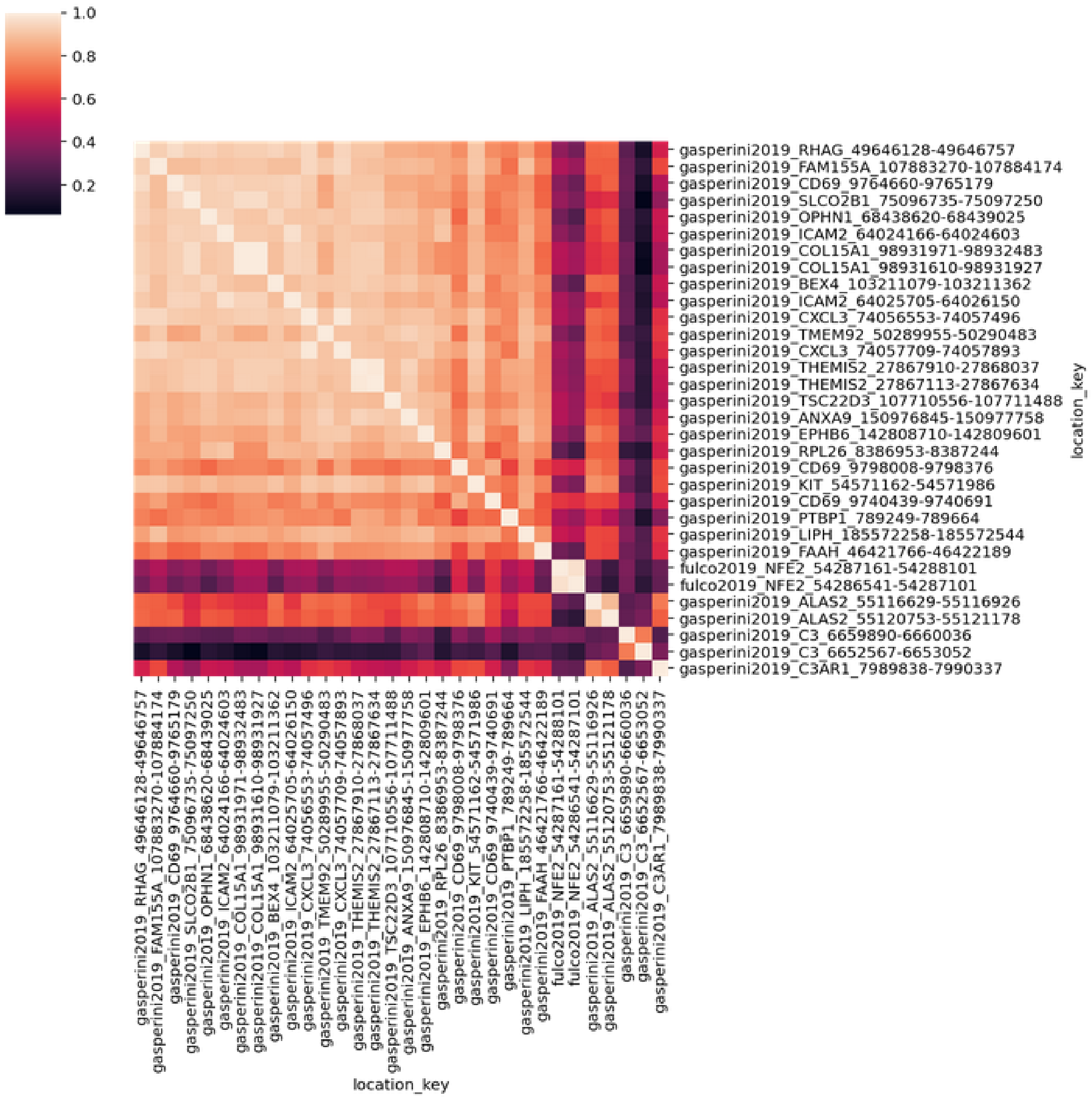


***Supplementary Figure 21**: Correlation of enhancer strengths estimated for different backgrounds. The backgrounds are sorted by the amount of variance explained by the enhancer strengths, i.e. the backgrounds at the very bottom are those where enhancers overall had little impact on expression at the promoter. We see that, if enhancers matter, their effects are highly consistent across backgrounds.*

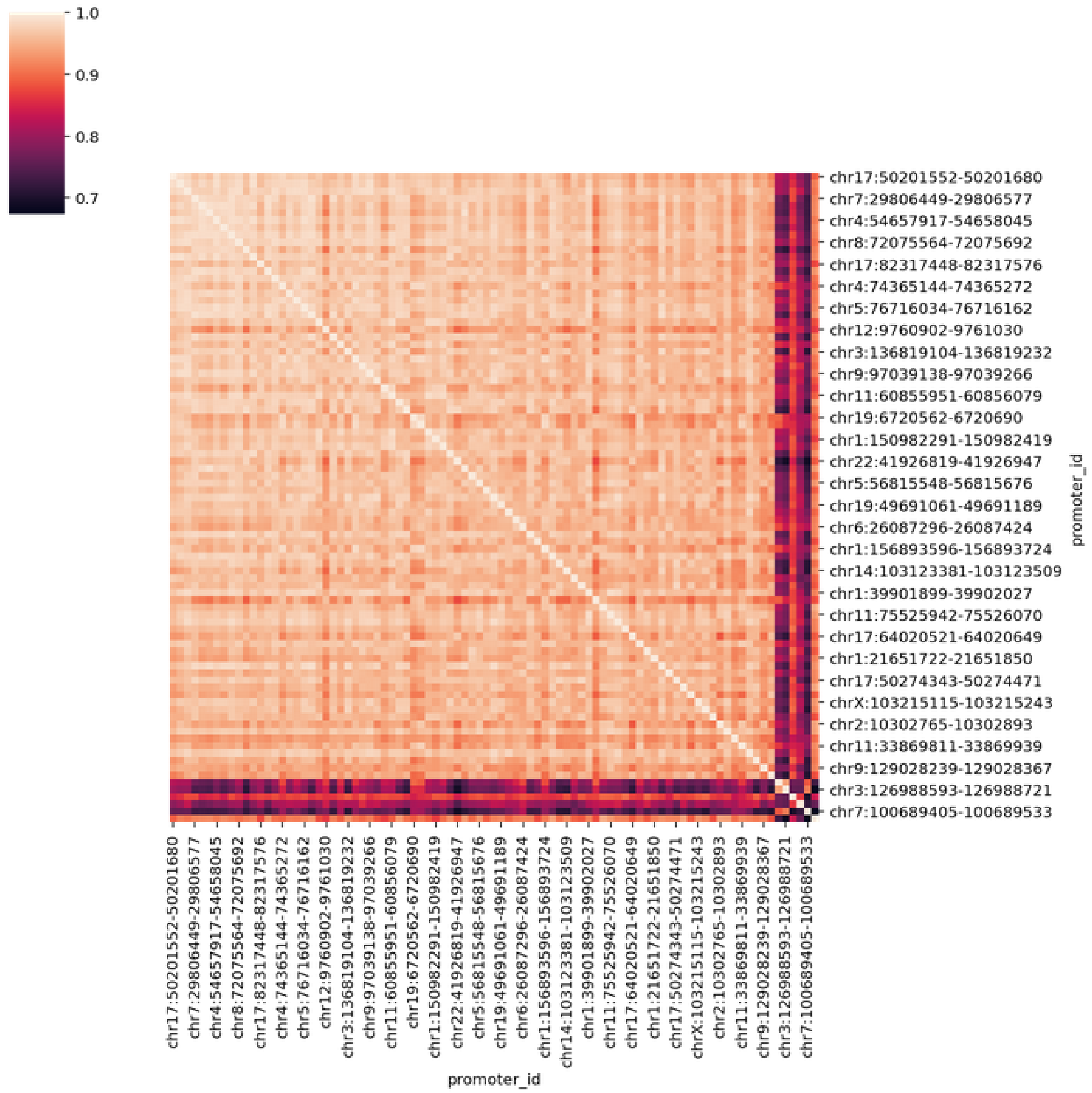


***Supplementary Figure 22**: Correlation of predicted enhancer strengths between promoters. The promoters are sorted by the amount of variance explained by the estimated enhancer strengths, i.e. the promoters at the very bottom are those which seemed to be mostly unaffected by enhancers. We see that, if enhancers matter, their effects are highly consistent across promoters.*
